# Structures of pUG-fold RNA bound to DNMT1 reveal a mechanism for RNA-mediated epigenetic regulation

**DOI:** 10.64898/2026.09.01.748594

**Authors:** Jessica J. Song, Thomas R. Cech, Vignesh Kasinath

## Abstract

Many chromatin-associated proteins have been found to bind RNA as a means of epigenetic regulation. Specifically, DNA methyltransferase 1 (DNMT1), which maintains cytosine methylation at CpG dinucleotides, is inhibited by RNA at transcribed DNA loci in cells. However, the mechanisms by which RNA binds DNMT1 and inhibits its activity remain unknown. Here, we determine a series of cryogenic electron microscopy (cryo-EM) structures of human DNMT1 bound to pUG-fold RNA, a non-canonical G-quadruplex previously observed to inhibit activity, revealing two distinct RNA-binding modes. The pUG-fold RNA binds the surface of DNMT1 in its autoinhibited conformation across a positively charged surface between the methyltransferase domain and the CXXC domain, and it binds directly in the active site of an open DNMT1 conformation. RNA binding is sterically incompatible with substrate DNA engagement in both states. Our 2.5 Å structure captures the intricate network of hydrogen bonds and electrostatic interactions between amino acids in the methyltransferase domain and the tetrad layers of pUG-fold RNA. Metadynamics molecular dynamics simulations provide an orthogonal view of the conformational landscape of DNMT1, revealing the two distinct RNA-binding modes. Furthermore, our analysis of published DNMT1 RIP-seq and eCLIP-seq data confirms that DNMT1-interacting RNAs in cells exhibit a strong propensity to form non-canonical G-quadruplex RNA structures. Collectively, our study provides the first structural basis for pUG-fold RNA recognition by a protein and illustrates how cryo-EM and AI-based methods for protein and RNA structure prediction synergize to inform the mechanism of RNA-mediated regulation of DNMT1.

## Introduction

One of the most important epigenetic modifications in eukaryotes is the methylation at position 5 of cytosine (5mC) in the context of CpG dinucleotides. DNA methylation is essential for transcriptional repression at CpG-rich promoters, imprinted genes, inactivated X chromosomes, transposons, and pericentromeric satellite DNA repeats^1,2^. Approximately 70- 80% of CpGs in mammalian genomes are methylated, with subtle differences between tissues^3^. DNA methyltransferases (DNMTs) are a class of enzymes that establish and maintain this DNA methylation. DNMT3A and DNMT3B work with DNMT3L to *de novo* methylate the genome during development, whereas DNMT1 is the only enzyme responsible for DNA methylation maintenance during DNA replication^1,2^. Regulation of DNMTs is crucial for faithfully propagating methylation patterns from parent cell to daughter cell.

Human DNMT1 is the largest DNMT with a large N-terminal regulatory region and a C-terminal catalytic region (Fig. 1a). The highly conserved methyltransferase (MTase) domain forms the core of the protein. The MTase domain utilizes S-adenosylmethionine (SAM) as a methyl donor to symmetrically methylate hemi-methylated DNA, in which one strand contains 5mC marks and the other is unmethylated. The first 350 amino acids of DNMT1 are largely unstructured. This region has been implicated in interactions with various binding partners, such as DNMT1-associated protein 1 (DMAP1) and proliferating cell nuclear antigen (PCNA)^4,5^, and they appear to alleviate non-productive binding of DNMT1 to chromatin^6^. The replication foci targeting sequence (RFTS) domain forms the top lobe of the protein, as seen by X-ray crystallography, and is responsible for recruitment to replication forks^7,8^. The RFTS domain can anchor to a cleft in the methyltransferase domain, thereby blocking DNA binding and methyltransferase activity^7^. Alternatively, the RFTS domain interacts with ubiquitinated histone H3 tails to relieve autoinhibition^9–12^. CXXC is a zinc finger domain that binds to unmethylated DNA^13^. A region between the CXXC domain and the N-terminal BAH domain, termed the autoinhibitory linker, contains a helix that bridges the RFTS and catalytic domains and strengthens this autoinhibitory interaction^7,13,14^. Together, the RFTS and CXXC domains regulate the conformational state of DNMT1 and subsequent activity.

**Fig. 1:**
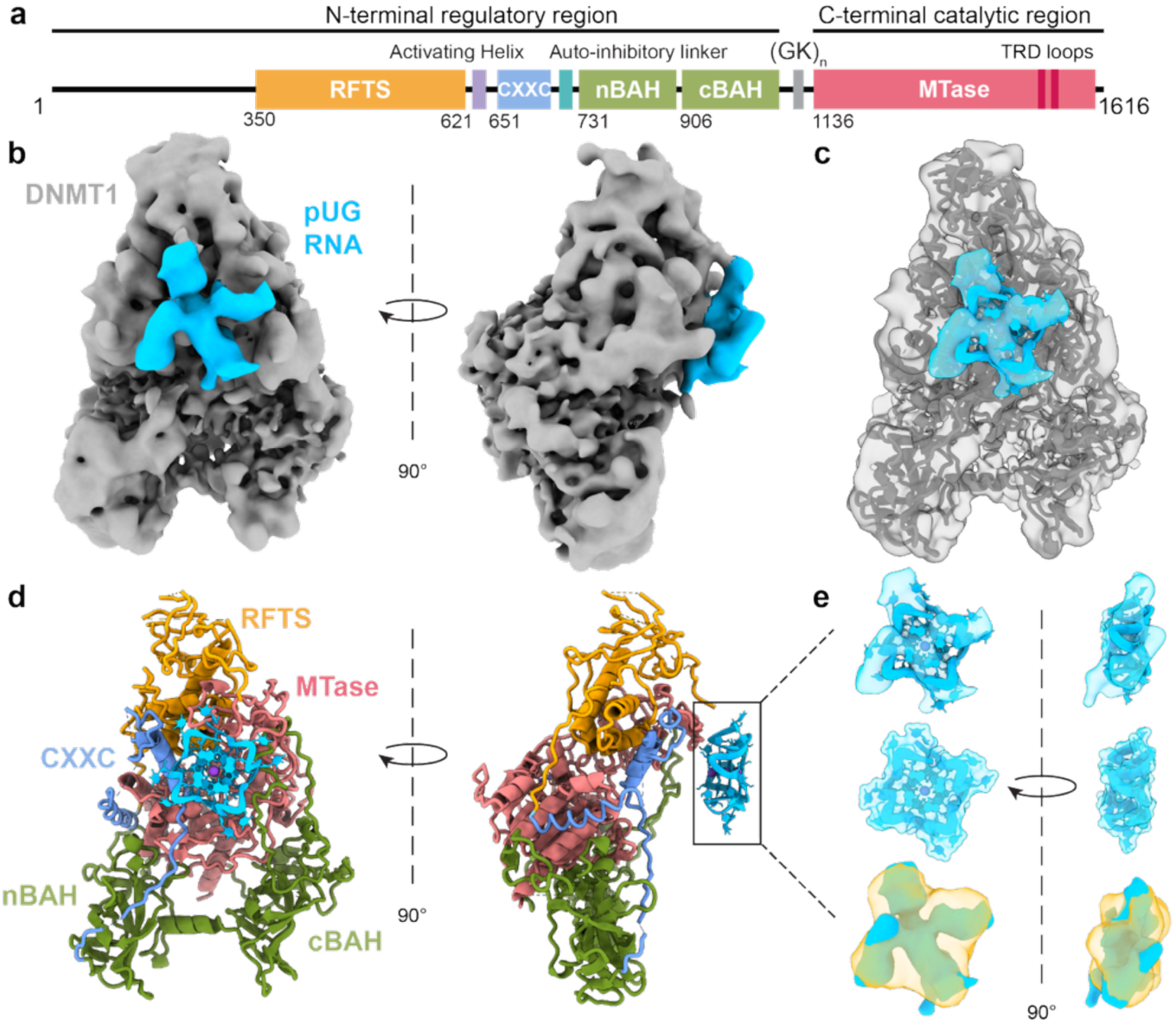
**pUG-fold RNA binds to the surface of DNMT1 to block DNA binding**. **a,** Domain organization of human DNMT1. **b,** Cryo-EM reconstruction of the full-length DNMT1 (gray) and (GU)_20_ pUG-fold RNA (blue) complex. **c,** Atomic models of DNMT1 and pUG-fold RNA fit into the cryo-EM map (PDB: 38DB). **d,** Atomic models of the DNMT1-pUG-fold RNA complex with RNA in blue, RFTS domain in orange, n-BAH and c-BAH in olive green, CXXC domain in cornflower blue, and catalytic MTAse domain in light coral. **e,** Superposition of the pUG-fold RNA NMR structure (PDB: 8TNS) in the cryo-EM density map (top), the NMR structure rendered as a surface (middle), and the surface rendered at 8 Å superimposed with the cryo-EM density (yellow).

DNMT1 lacks a canonical RNA recognition motif; however, like many other chromatin-associated proteins, it has been found to bind to and be regulated by RNA^15^. Di Ruscio et al. identified long noncoding RNA (lncRNA) ecCEBPA as a regulator of DNMT1 in cells and discovered that RNA inhibition of DNMT1 occurs genome-wide^16^. Other studies have found that DNMT1 directly interacts with the lncRNA DACOR1, mir-155-5p, and nascent mRNAs, with most of these interactions resulting in inhibition of DNA methyltransferase activity^17–19^. Additionally, a recent study has identified many more RNAs that associate with DNMT1, including the tumorigenesis-associated lncRNA LINC00339 and the nucleolar phosphoprotein chaperone NPM1 RN^20^. Our previous work has shown that DNMT1 binds strongly to both canonical G-quadruplex RNA and the non-canonical form consisting of GU repeats, termed poly(UG)-fold (pUG-fold)^21^. The pUG-fold requires 12 GU repeats and potassium ions to fold but can tolerate some sequence variability^22,23^. This unique RNA fold consists of three G quartets and one U quartet stacked on top of one another, with the remaining uridines flipped out^22,24^. Both canonical RNA G-quadruplexes and the pUG-fold inhibit DNMT1 activity, with the pUG-fold RNA being a stronger inhibitor *in vitro*^21^. It remains unknown how DNMT1 interacts with pUG- fold RNA and how engagement of pUG-fold RNA inhibits DNMT1 activity.

Here, we investigate how pUG-fold RNA binds DNMT1 and how RNA binding affects the conformational state of DNMT1. Using a series of cryo-EM structures of full-length and mutated DNMT1 constructs in complex with pUG-fold RNA, together with AlphaFold3 predictions, we show that the RNA can bind to DNMT1 in multiple locations. The RNA can bind either on the protein surface to block DNA binding or directly within the DNA-binding site in the methyltransferase domain. Metadynamics molecular dynamics simulations reveal how mutations in the CXXC and methyltransferase domains disrupt the RFTS/CXXC-mediated autoinhibition and facilitate RNA binding. Our study presents the first structure of pUG-fold RNA recognition by a protein, illustrating a functionally relevant mechanism for RNA-mediated regulation of DNMT1, in which RNA and DNA compete for DNMT1 binding.

## Results

### pUG-fold RNA binds to the surface of DNMT1

To determine the structural basis for pUG-fold RNA recognition by human DNMT1, we assembled recombinant human full-length DNMT1 with a pUG-fold-forming RNA *in vitro* and subjected the complex to cryo-EM structural analysis. We obtained a 3.6 Å structure of a 1:1 RNA:DNMT1 complex (Fig. 1b-d, Extended Data Fig. 1, 2a-d, Table 1). The overall architecture of full-length DNMT1 in the complex resembles previously reported crystal and cryo-EM structures of apo-DNMT1, in which the RFTS domain directly associates with the methyltransferase domain and the autoinhibitory linker. Similar to previous cryo-EM structures of DNMT1^14^, the unstructured N-terminus (aa 1-350) could not be resolved in our reconstruction of the DNMT1-RNA complex.

**Table 1.** Cryo-EM data collection, refinement, and validation statistics

| | Full-length<br>DNMT1+RNA | Mut3+CXXCmut<br>DNMT1-RNA | Δ1-619 DNMT1-RNA |
| --- | --- | --- | --- |
| PDB accession code | 38DB | 38EM | 38DT |
| EMDB accession code | EMD-78737 | EMD-78766 | EMD-78750 |
| Data collection and processing |  |  |  |
| Microscope | Krios 2 | Krios 2 | Krios 2 |
| Camera | Gatan energy<br>filtered K3 | Gatan energy<br>filtered K3 | Gatan energy<br>filtered K3 |
| Energy Filter | BioContinuum | BioContinuum | BioContinuum |
| Magnification | 130kx | 130kx | 130kx |
| Voltage (kV) | 300 | 300 | 300 |
| Electron exposure ($e^-/\text{\AA}^2$) | 50 | 50 | 50 |
| Defocus range ($\mu\text{m}$) | -0.6 to -2.0 | -0.8 to -2.5 | -0.8 to -2.5 |
| Pixel size ($\text{\AA}$) | 0.6488<br>(super res 0.3244) | 0.6485<br>(super res<br>0.32425) | 0.8257<br>(super res<br>0.41285) |
| Symmetry imposed | C1 | C1 | C1 |
| Initial particles images (no.) | 2,963,729 | 8,033,665 | 12,056,892 |
| Final particles images (no.) | 32,573 | 226,984 | 349,666 |
| Map resolution ($\text{\AA}$) | 3.6 | 2.8 | 2.5 |
| FSC threshold | 0.143 | 0.143 | 0.143 |
| Map resolution range (Å) | 4-10 | 3-5 | 2.2-3.5 |
| **Refinement** |  |  |  |
| Initial model used (PDB code) |  |  |  |
| Model resolution (Å) | 3.8 | 3.2 | 2.7 |
| FSC threshold | 0.5 | 0.5 | 0.5 |
| Map sharpening B factor (Å<sup>2</sup>) | 88 | 65 | 30 |
| Model composition |  |  |  |
| Nonhydrogen atoms | 8459 | 6879 | 7009 |
| Protein residues | 1133 | 839 | 825 |
| Nucleotide residues | 24 | 24 | 24 |
| Ligands | 8 | 6 | 6 |
| *B* factors (Å<sup>2</sup>) |  |  |  |
| Protein (mean) | 30.44 | 87.52 | 60.77 |
| Nucleotide (mean) | 176.29 | 179.66 | 67.51 |
| Ligand (mean) | 25.44 | 84.49 | 54.13 |
| R.m.s deviations |  |  |  |
| Bond lengths (Å) | 0.005 | 0.002 | 0.003 |
| Bond angles (°) | 0.899 | 0.492 | 0.574 |
| Validation |  |  |  |
| MolProbity score | 2.19 | 1.45 | 0.88 |
| Clash score | 6.12 | 4.82 | 1.18 |
| Poor rotamers (%) | 3.30 | 0.47 | 0.58 |
| Ramachandran plot |  |  |  |
| Favored (%) | 92.74 | 96.74 | 97.80 |
| Allowed (%) | 6.64 | 3.26 | 2.09 |
| Disallowed (%) | 0.63 | 0 | 0.12 |

We resolved an additional density on the surface of the methyltransferase domain, which we assigned to RNA, as it had not been observed in previous structures of apo-DNMT1^7,14^. This RNA density is approximately the same shape and size as the NMR structure of the pUG-fold RNA^24^ (Fig. 1e). Our structure predicts that RNA in this position would clash with DNA binding in the active site. To assess the potential overlap between the pUG-fold RNA and DNA, we rigid-body docked a hemi-methylated 19-mer DNA in the catalytic site (PDB 4DA4) into the cryo-EM density map (Extended Data Fig. 3a)^13^. The RNA density blocked the DNA exit site in the catalytic domain. We further modeled a 31-mer DNA (PDB 7Q94) to illustrate the overlap between binding sites (Fig. 2a). We also rigid-body docked unmethylated 12-mer DNA bound to the CXXC domain (PDB 3PTA) into the density map (Extended Data Fig. 3b) and found that the unmethylated DNA also sterically clashes with the RNA^25^. To validate these structural analyses, we carried out biochemical binding competition assays, which confirmed that RNA and DNA cannot simultaneously occupy DNMT1 (Extended Data Fig. 3g). Therefore, RNA binding directly in front of the catalytic methyltransferase domain results in a steric clash with the DNA substrate binding at the active site.

**Fig. 2:**
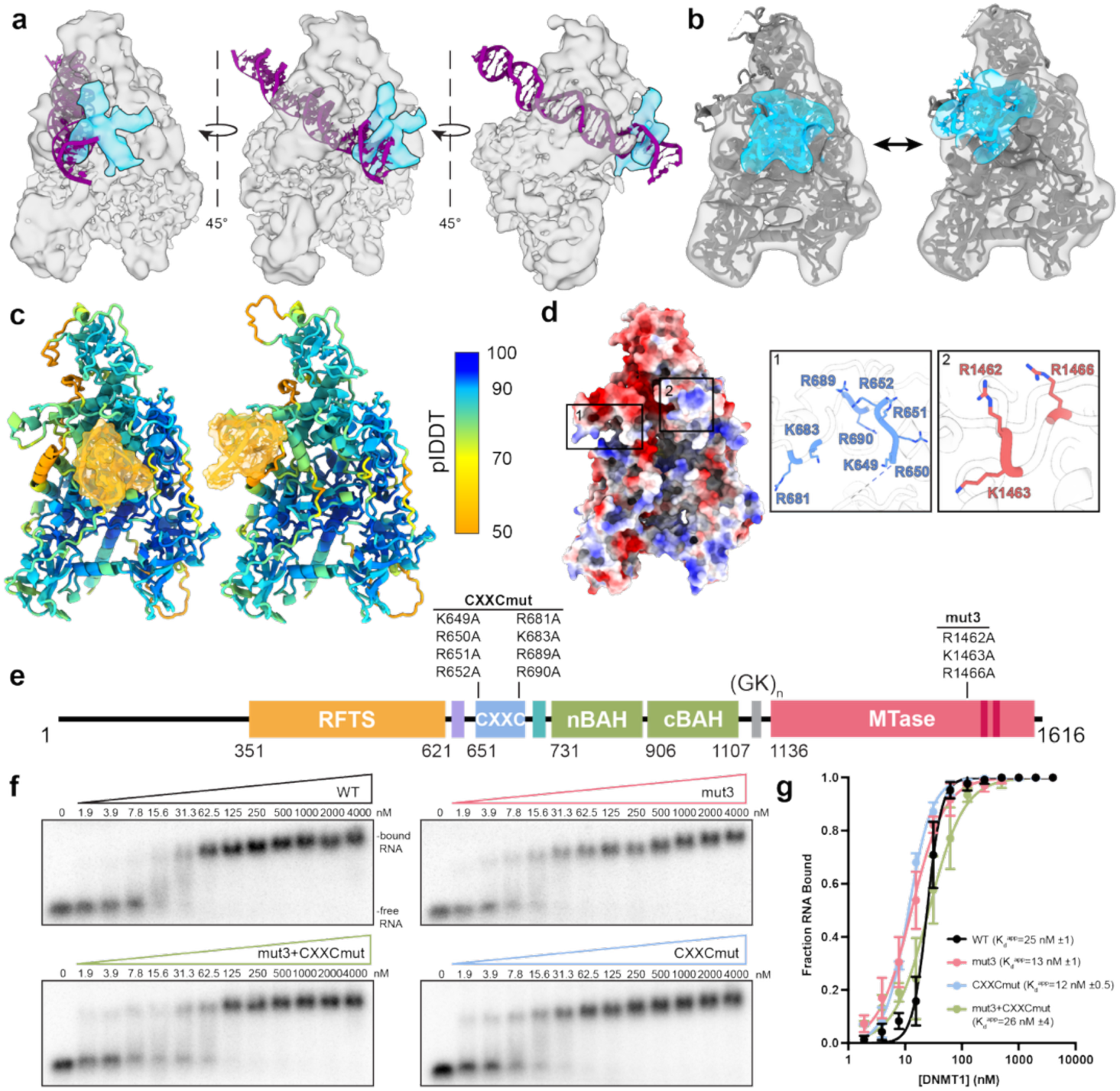
Mutations in the RNA-binding surface do not affect RNA binding. a, Rigid body docking of DNA (purple; PDB: 7Q94) in the active site of DNMT1 shows the clash between pUG-fold RNA and DNA. **b,** 3D variability analysis shows a shift in RNA density (blue) across the surface of DNMT1 (gray), away from the methyltransferase and CXXC domains. **c,** AlphaFold3 predictions of full length DNMT1 and TERRA RNA ([UUAGGG]_4_, K^+^). Residues 1- 350 omitted for clarity. Models colored by predicted Local Distance Difference Test (plDDT). Predicted aligned error (PAE) plots for both models in Supplemental Figure 1. **d,** Electrostatic surface of DNMT1 (PDB: 4WXX) and inset of (1) CXXCmut residues and (2) mut3 residues. **e,** Domain schematic of mut3+CXXCmut DNMT1 with point mutations. **f,** Representative EMSAs of trace amounts of (GU)_20_ RNA and increasing amounts of indicated DNMT1 construct. **g,** Binding curves of DNMT1 constructs and (GU)_20_ RNA from panel F. Points are mean values and error bars represent standard deviations for 3 independent measurements.

Three-dimensional variability analysis (3DVA) of the full-length DNMT1-pUG-fold RNA cryo-EM dataset revealed continuous conformational heterogeneity in which the RNA density moves along a surface between the methyltransferase domain and the CXXC domain (Fig. 2b). Intrigued by RNA binding broadly across this DNMT1 surface, we used AlphaFold3 to predict the different binding states between DNMT1 and TERRA RNA^26^. We used TERRA RNA [UUAGGG)_4_], the human telomeric repeat RNA, because AlphaFold3 was unable to fold (GU)_20_ RNA into a pUG-fold but could fold TERRA into a G-quadruplex. AlphaFold3 predicted two distinct RNA-binding sites – the primary interface identified by cryo-EM in front of the methyltransferase domain and a secondary interface proximal to the CXXC domain, as observed by our 3DVA (Fig. 2c, Supplemental Fig. 1). These AlphaFold3 predictions are low confidence but are validated by our cryo-EM structural analysis. Together, these orthogonal methods demonstrate that G-quadruplex-forming RNAs may engage DNMT1 along an extended interface to sterically block DNA binding.

## Mutations of the RNA-binding surface do not affect RNA binding

In an attempt to validate the amino acid residues involved in pUG-fold RNA binding, we generated mutations of positively charged lysine and arginine residues (referred to as mut3: R1462A K1463A R1466A) on the surface loop of the methyltransferase domain identified from our cryo-EM structure (Fig. 2d-e). However, to our surprise, electrophoretic mobility shift assays (EMSA) revealed mut3 bound more tightly to (GU)_20_ RNA compared to wild-type (WT) (K_d_^app^ = 13 ± 1 nM [mut3] vs K_d_^app^ = 25 ± 1 nM [WT]) and no significant difference in hemi-methylated DNA binding compared to that of wild type (K_d_^app^ = 150 ± 9 nM [mut3] vs K_d_^app^ = 159 ± 42 nM [WT]) (Fig. 2f-g, Extended Data Fig. 3e-f, Table 2).

**Table 2.**
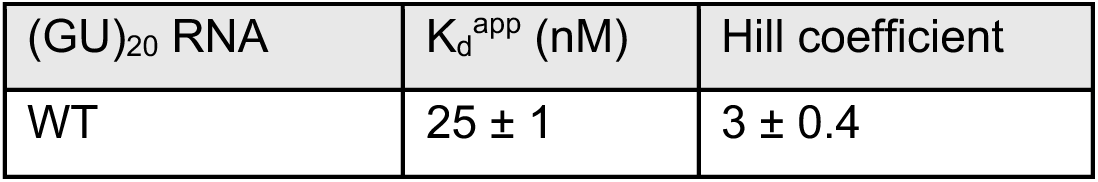
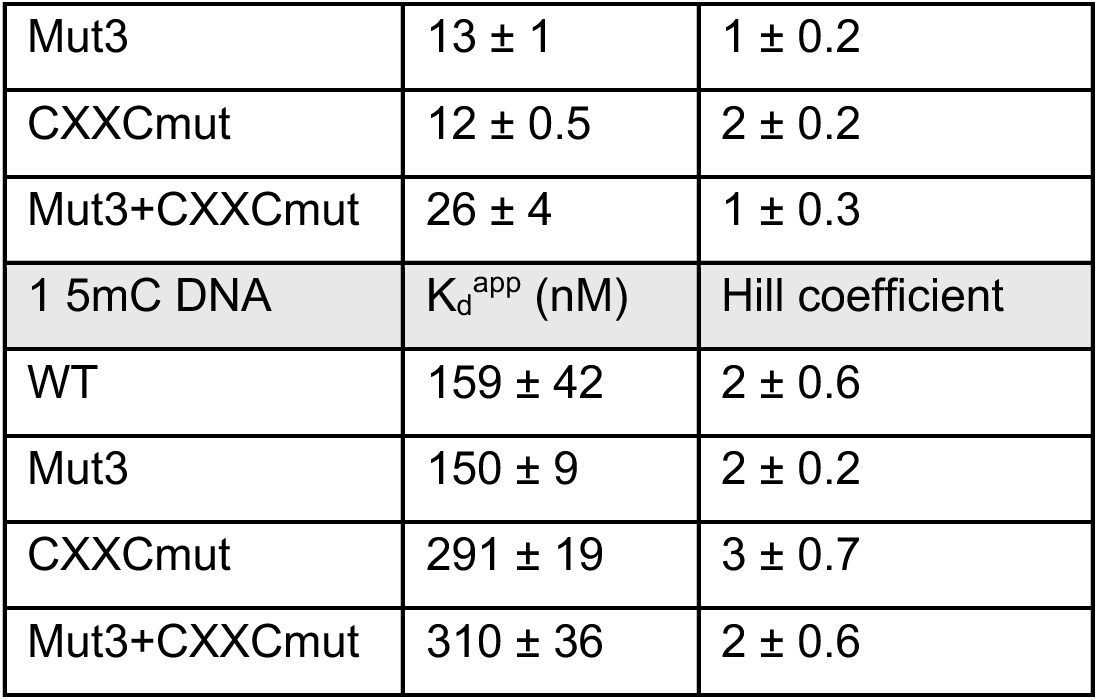
Equilibrium constants of (GU)_20_ RNA and 1 5mC DNA binding to DNMT1 constructs

We then hypothesized that charge disruption of residues in the methyltransferase domain resulted in stabilized RNA binding at the CXXC domain, as observed in 3DVA and AlphaFold3 predictions (Fig. 2b-c). To test this, we generated charge-neutralization mutations in the CXXC domain predicted to be involved in RNA binding by AlphaFold3 and previously implicated in unmethylated DNA binding (referred to as CXXCmut: K649A R650A R651A R652A R681A K683A R689A R690A)^13^, as well as a combination mutant targeting both the mut3 and CXXCmut residues (referred to as mut3+CXXCmut) (Fig. 2d-e). We intentionally did not mutate residues involved in hemi-methylated DNA coordination (e.g., M1232, R1234, K1535) or methyltransferase activity (e.g., C1226, E1266, R1310, R1312) to avoid disrupting enzymatic activity. AlphaFold3 predictions of DNMT1 mut3, CXXCmut, and mut3+CXXCmut were nearly identical to WT DNMT1 (predicted template modeling score (pTM) = 0.75), confirming that these mutations did not affect the overall fold (Supplemental Fig. 2). EMSA binding assays again showed a two-fold increase in CXXCmut binding affinity for (GU)_20_ RNA (K_d_^app^ = 12 ± 0.5 nM), but a nearly two-fold decrease in binding affinity for hemi-methylated DNA (K_d_^app^ = 291 ± 19 nM). Surprisingly, the binding affinity of the mut3+CXXCmut mutant for (GU)_20_ RNA was the same as WT (K_d_^app^ = 26 ± 4 nM) despite mutation of two discrete RNA-binding sites, while the binding affinity for hemi-methylated DNA followed the same trend as the CXXCmut, with a twofold decrease in binding affinity (K_d_^app^ = 310 ± 36 nM) (Fig. 2f-g, Extended Data Fig. 3e-f, Table 2). The weakened binding affinity for DNA in the CXXCmut and combination mutant is likely due to the presence of only one 5mCpG site on the DNA substrate and the CXXC domain’s known role in binding unmethylated DNA. However, the unchanged (GU)_20_ RNA binding affinity in the combination mutant suggests an additional RNA-binding site on DNMT1.

Furthermore, in binding competition assays in which DNMT1 was pre-bound to hemi-methylated DNA and increasing amounts of (GU)_20_ RNA were added, RNA displaced the DNA from WT DNMT1 (IC_50_ ∼ 125 nM) and less effectively with the mut3+CXXCmut protein (IC_50_ ∼500 nM). The RNA is likely less effective at displacing the DNA in the combination mutant because protein surface binding is disrupted, suggesting a second binding site where the RNA and DNA directly compete for the same site. When DNMT1 was pre-bound to (GU)_20_ RNA, and increasing amounts of hemi-methylated DNA were added, the DNA did not displace the RNA from either the WT or combination mutant during the 60-minute incubation period of these experiments (Extended Data Fig. 3g, Supplemental Fig. 3). The lack of displacement can be explained by the minor population of DNMT1 in a conformation competent for DNA binding (see Fig. 5 analysis).

These findings suggest that pUG-fold RNA binding by DNMT1 does not depend on a single or a few critical electrostatic contact points but may instead be mediated by a distributed network of dynamic interactions across the protein surface and/or potentially at a second discrete site. This manner of binding would confer robustness against single-point mutagenesis of amino acids identified as RNA-binding by cryo-EM, as the RNA will likely shift to another site on the protein. This interpretation is also consistent with the variety of RNA sequences and structures DNMT1 can bind^16,21^. Such limited sequence specificity is a common theme among chromatin-associated proteins that lack a canonical RNA-binding domain, including CTCF, PRC2, and YY1^27–29^.

## pUG-fold RNA has an additional binding site on DNMT1

Because mutations in the catalytic and CXXC domains did not disrupt RNA binding, we sought to understand how pUG-fold RNA engages the combination mutant mut3+CXXCmut DNMT1. Since hemi-methylated DNA engagement with the catalytic site of DNMT1 requires dislocation of the CXXC and RFTS domains, we hypothesized that mutations in the CXXC domain contribute to conformational flexibility that exposes the catalytic site for pUG-fold RNA binding. To test this, we assembled a complex of the mut3+CXXCmut DNMT1 protein and (GU)_20_ RNA and utilized cryo-EM to determine additional pUG-fold binding sites on DNMT1. Structural analysis yielded two conformations of DNMT1: an RFTS and CXXC disengaged state (referred to as open-bound state) and a closed, autoinhibitory state (referred to as closed-unbound state), where “bound” and “unbound” refer to RNA binding (Extended Data Fig. 4c).

In the open-bound state, the RFTS and CXXC domains and the autoinhibitory domain could not be resolved (Fig. 3a-c, Extended Data Fig. 4, 5a, Table 1), resulting in an open conformation of the mutant DNMT1 that broadly resembles previously reported structures of WT DNMT1 bound to unmethylated DNA^13,30^ (Supplemental Fig. 4a-d). We unambiguously resolved the pUG-fold RNA in the catalytic site of DNMT1 at 4-5 Å resolution (Fig. 3a, b). Closer structural analysis revealed that the pUG-fold RNA pushes the catalytic loop (residues 1224- 1238) down towards the n-BAH domain, approximately 16 Å relative to the active conformation (Extended Data Fig. 5e). Additionally, the α-helix after the catalytic loop spanning residues 1239-1250), which adopts a straight conformation in the active state (PDB: 4DA4; hDNMT1 aa 729-1602 bound to hemi-methylated DNA), has a kink similar to that observed in the inactive states (PDB: 3PTA – hDNMT1 aa 646-1600 bound to unmethylated DNA and PDB: 4WXX –apo-, autoinhibited hDNMT1 aa 351-1600) (Extended Data Fig. 5e)^13,25^. While we were able to capture the conformation of DNMT1, the RNA resolution was insufficient to determine atomic-level interactions between DNMT1 and the pUG-fold RNA.

**Fig. 3:**
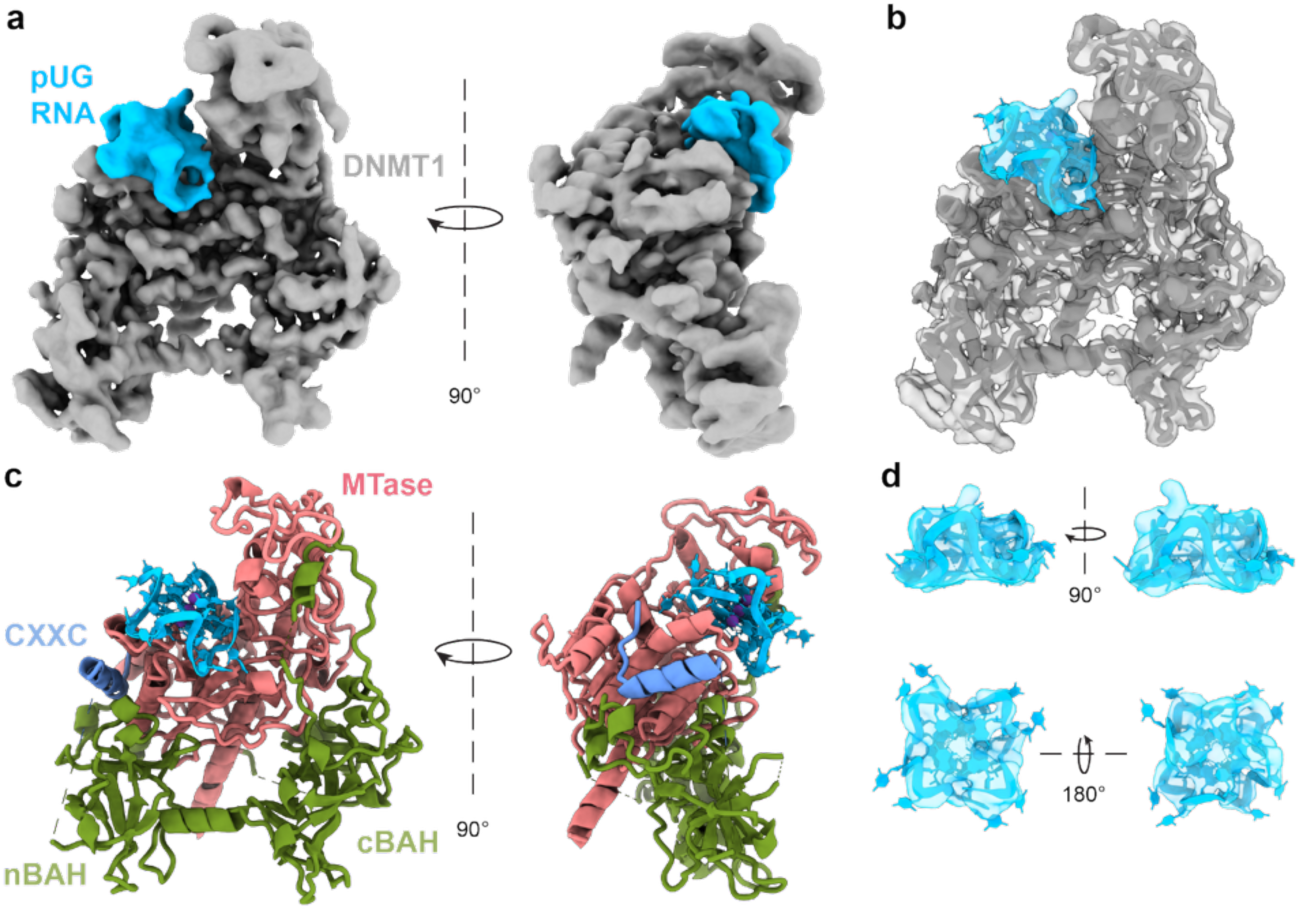
pUG RNA has a second binding site on DNMT1. a, Cryo-EM density map of mut3+CXXCmut DNMT1 (gray) bound to pUG-fold RNA (blue). **b,** Atomic models mut3+CXXCmut DNMT1 (gray) bound to pUG-fold RNA docked into the cryo-EM map (PDB: 38EM). **c,** Atomic models of mut3+CXXCmut DNMT1-(GU)_20_ colored by DNMT1 domain. **d,** Atomic model of the pUG RNA docked into the cryo-EM map.

In the closed-unbound state, the predominant class in the mut3+CXXCmut DNMT1- (GU)_20_ RNA dataset, the mutant DNMT1 protein adopts an overall conformation indistinguishable from the wild-type full-length DNMT1 structures and our own structure (Extended Data Fig. 4c, Fig. 1, Supplemental Fig. 4e-f)^7,14^. The RFTS domain and autoinhibitory linker associate with the catalytic domain to render the protein in an autoinhibited conformation. We did not observe any interpretable RNA density on the surface of the protein. This is expected due to the mut3+CXXCmut mutations, which were designed to disrupt surface engagement of the RNA. The absence of RNA in this cryo-EM map indicates that RFTS disengagement from the catalytic domain is required for RNA binding in the catalytic site. Collectively, these data reveal how conformational equilibrium of DNMT1 governs RNA accessibility to the substrate site and suggest that factors or mutations that promote RFTS disengagement and relief of autoinhibition would sensitize DNMT1 to RNA engagement and subsequent inhibition.

## RNA binds to the catalytic site of DNMT1 to render the enzyme inactive

To accurately map interactions between the pUG-fold RNA and the open conformation of the protein, we deleted the unstructured N-terminus and RFTS domain of (Δ1-619) (Fig. 4a). We then assembled this truncated protein with (GU)_20_ RNA and subjected the complex to cryo-EM structural analysis. We hypothesized that, in the absence of the unstructured N-terminus and the RFTS domain, the catalytic site would be exposed to pUG-fold RNA binding.

**Fig. 4:**
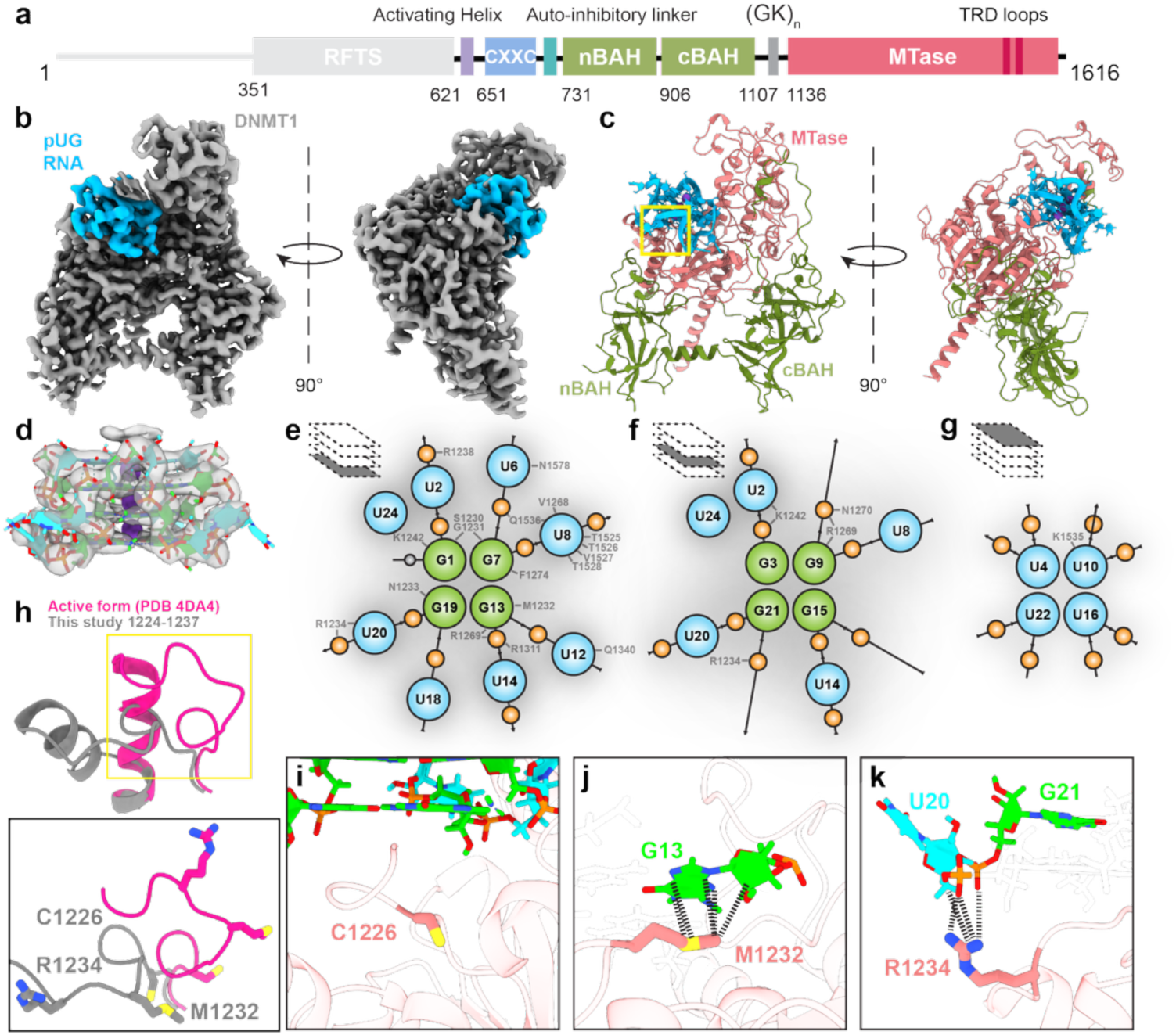
pUG-fold RNA binds to the catalytic site of DNMT1. a, Domain schematic of Δ1-619 DNMT1. **b,** Cryo-EM density map of Δ1-619 DNMT1 (gray) bound to pUG-fold RNA (blue). **c,** Atomic models of Δ1-619 DNMT1 bound to pUG-fold RNA, colored by DNMT1 domain. Yellow box represents zoomed-in view in H (PDB: 38DT). **d,** Atomic model of fit into the cryo- EM density. Potassium ions are represented as purple spheres, guanine bases are lime, uracil bases are cyan, and phosphates are orange. **e,** Schematic of interactions between Δ1-619 DNMT1 and the G1 quartet of pUG-fold RNA. **f,** Schematic of interactions between Δ1-619 DNMT1 and the G3 quartet of pUG-fold RNA. **g,** Schematic of interaction between Δ1-619 DNMT1 and the U4 quartet of pUG-fold RNA. Hydrogen bonds, electrostatic interactions, and Van der Waals interactions shown. Phosphates shown as orange spheres. **h,** Comparison of the catalytic loop and adjacent α-helix in the active form DNMT1 bound to hemi-methylated DNA (deep pink, PDB: 4DA4) and this study (gray). Inset below specifically compares the conformation and orientation of residues C1226, M1232, and R1234 to the corresponding residues in the active form. The catalytic cysteine residue (C1226) is positioned away from the DNA binding site. **i,** M1232 forms van der Waals contacts with the base and sugar of G13. **j,** R1234 forms electrostatic contacts with the phosphate groups of nucleotides U20 and G21. **k,** R1234 forms electrostatic contacts with the phosphate groups of nucleotides U20 and G21.

We determined the cryo-EM structure of the truncated DNMT1 (Δ1-619) in complex with pUG-fold RNA to an overall resolution of 2.5 Å (Fig. 4b-c and Extended Data Fig. 6, 7a, Table 1). At this resolution, the phosphodiester backbone of the pUG-fold RNA could be unambiguously traced within the cryo-EM density (Fig. 4d). The four distinct layers of the pUG-fold, comprised of three G-quartets and one U-quartet, were easily visible. Electron density corresponding to a bound K^+^ ion was seen between each planar G quartet and between the top G-quartet (G5 quartet) and the U-quartet (Fig. 4d, Supplemental Fig. 7f). The flipped-out uracil nucleotides, unique to the pUG-fold, were seen at lower volume thresholds (Supplemental Fig. 7d).

The high-resolution map allowed us to identify a network of 6 hydrogen bonds, 14 electrostatic interactions, and nearly 50 van der Waals contacts between the RNA and DNMT1 (Fig. 4e-g, Extended Data Fig. 8, Table 3). The RNA is tightly nestled into the catalytic site, at the same location as the RNA in complex with mut3+CXXCmut DNMT1 (Fig. 4b-c, Extended Data Fig. 7f). Similarly, the pUG-fold RNA pushes the catalytic loop (residues 1224-1238) down towards the n-BAH domain and the α-helix after the catalytic loop is kinked (Fig. 4h, Extended Data Fig. 7g). The catalytic cysteine C1226 is pointed down away from the RNA, in a manner incompatible with productive methylation (Fig. 4h-i). In the previous structure of hemi-methylated DNA-bound DNMT1 (PDB:4DA4), the side chain of M1232 intercalates into the DNA to occupy the space from the flipped-out target cytosine, and R1234 further binds the DNA (Fig. 4h). Mutations of M1232 and R1234 to alanine result in nearly 8-fold and 4-fold decreases in activity, respectively^25^. Here, the sulfur atom of the M1232 side chain packs against the G13 nucleotide ring face within the G1 quartet, while also forming a chalcogen contact with the same base (Fig. 4j). R1234 forms electrostatic contacts with a phosphate group of G21, one of the guanosine nucleotides in the G3 quartet, and a phosphate group of U20, a flipped-out uracil base (Fig. 4k). Therefore, RNA engagement of residues critical for binding DNA in the active site contributes to the pUG-fold-mediated mechanism of inhibition.

**Table 3.** Interactions between DNMT1 (Δ1-619) and (GU)_20_ RNA

| Donor | Donor atom | Acceptor | Acceptor atom | D...A (Å) | Est. D-H...A (°) | Classification |
| --- | --- | --- | --- | --- | --- | --- |
| ASN 1578 | ND2 | U6 | O4 | 2.89 | ~92° | Hydrogen bond |
| ARG 1269 | NH1 | G9 | OP1 | 2.93 | ~98° | Hydrogen bond |
| THR 1528 | N | U8 | O4 | 3.08 | ~139° | Moderate Hydrogen bond |
| ASN 1270 | ND2 | G9 | OP2 | 3.10 | ~150° | Moderate Hydrogen bond |
| LYS 1242 | NZ | G1 | O2' | 3.41 | ~136° | Weak Hydrogen bond |
| ARG 1269 | NH1 | G9 | O5' | 3.45 | ~139° | Weak Hydrogen bond |
| GLY 1231 | N | G1 | O6 | 3.45 | ~57° | Probable H-bond |
| ARG 1269 | NE | G9 | OP1 | 3.04 | ~76° | Probable H-bond |
| ARG 1269 | NH1 | G13 | OP1 | 2.94 | ~88° | Probable H-bond |
| ARG 1269 | NH2 | G13 | OP1 | 3.03 | ~84° | Probable H-bond |
| ARG 1269 | NH1 | G13 | OP2 | 3.95 | — | Electrostatic |
| ARG 1269 | NH1 | G9 | OP2 | 4.17 | — | Electrostatic |
| ARG 1269 | NE | G13 | OP1 | 4.21 | — | Electrostatic |
| LYS 1242 | NZ | U2 | OP2 | 4.32 | — | Electrostatic |
| LYS 1242 | NZ | G3 | OP2 | 4.37 | — | Electrostatic |
| ARG 1234 | NH1 | U20 | OP2 | 4.45 | — | Electrostatic |
| ARG 1238 | NE | U2 | OP2 | 4.45 | — | Electrostatic |
| ARG 1269 | NE | G9 | OP2 | 4.65 | — | Electrostatic |
| ARG 1269 | NH2 | G9 | OP1 | 4.66 | — | Electrostatic |
| ARG 1311 | NH1 | G13 | OP1 | 4.71 | — | Electrostatic |
| ARG 1234 | NH2 | U20 | OP2 | 4.78 | — | Electrostatic |
| ARG 1311 | NH2 | G13 | OP1 | 4.78 | — | Electrostatic |
| ARG 1234 | NH2 | G21 | OP2 | 4.88 | — | Electrostatic |
| ARG 1269 | NH2 | G13 | OP2 | 4.99 | — | Electrostatic |
| PHE 1274 | ring | G7 | purine ring | 4.92 | 73.3° | T-shaped aromatic |
| MET 1232 | SD | G13 | ring C5/C6 | 3.89 | — | vdW packing |
| ASN 1578 | OD1 | U6 | O4 | 2.81 | — | Van der Waals (O⋯O) |
| GLN 1340 | OE1 | U12 | O2 | 2.95 | — | Van der Waals (O⋯O) |
| MET 1232 | SD | G13 | N1 | 3.92 | — | Chalcogen contact (S⋯N) |
| GLY 1231 | CA | G1 | O6 | 2.92 | — | Van der Waals |
| VAL 1527 | CA | U8 | O4 | 3.03 | — | Van der Waals |
| ARG 1269 | CD | G9 | OP1 | 3.04 | — | Van der Waals |
| ARG 1269 | CZ | G13 | OP1 | 3.18 | — | Van der Waals |
| ASN 1578 | CG | U6 | O4 | 3.23 | — | Van der Waals |
| SER 1230 | CB | G7 | N2 | 3.35 | — | Van der Waals |
| ARG 1269 | CZ | G9 | OP1 | 3.38 | — | Van der Waals |
| SER 1230 | O | G1 | C6 | 3.40 | — | Van der Waals |
| GLY 1231 | CA | G7 | O6 | 3.40 | — | Van der Waals |
| SER 1230 | CB | G7 | C2 | 3.41 | — | Van der Waals |
| SER 1230 | O | G1 | O6 | 3.42 | — | Van der Waals |
| ASN 1233 | CG | G19 | N7 | 3.43 | — | Van der Waals |
| ASN 1233 | CB | G19 | N7 | 3.43 | — | Van der Waals |
| ASN 1233 | CB | G19 | C8 | 3.44 | — | Van der Waals |
| GLY 1231 | N | G1 | O6 | 3.45 | — | Van der Waals |
| THR 1525 | O | U8 | C5 | 3.49 | — | Van der Waals |
| ASN 1233 | CB | G19 | C5 | 3.53 | — | Van der Waals |
| SER 1230 | O | G1 | C5 | 3.54 | — | Van der Waals |
| VAL 1527 | C | U8 | O4 | 3.56 | — | Van der Waals |
| ASN 1233 | CB | G19 | N9 | 3.60 | — | Van der Waals |
| ASN 1233 | CB | G19 | O2' | 3.61 | — | Van der Waals |
| LYS 1242 | NZ | U2 | C5' | 3.63 | — | Van der Waals |
| MET 1232 | CE | G13 | N3 | 3.63 | — | Van der Waals |
| VAL 1268 | CG1 | U8 | C5' | 3.64 | — | Van der Waals |
| GLY 1231 | CA | G1 | C6 | 3.65 | — | Van der Waals |
| THR 1526 | O | U8 | C4 | 3.65 | — | Van der Waals |
| VAL 1527 | CG1 | U8 | O4 | 3.65 | — | Van der Waals |
| ASN 1233 | CB | G19 | C4 | 3.65 | — | Van der Waals |
| LYS 1535 | CB | U10 | O2' | 3.67 | — | Van der Waals |
| GLN 1340 | CD | U12 | O2 | 3.69 | — | Van der Waals |
| SER 1230 | C | G1 | O6 | 3.69 | — | Van der Waals |
| ARG 1269 | NH1 | G9 | P | 3.70 | — | Van der Waals |
| GLN 1536 | CG | U8 | C5 | 3.72 | — | Van der Waals |
| VAL 1527 | CB | U8 | O4 | 3.72 | — | Van der Waals |
| ARG 1234 | NH2 | U20 | C5' | 3.75 | — | Van der Waals |
| ASN 1233 | CG | G19 | C8 | 3.78 | — | Van der Waals |
| MET 1232 | CE | G13 | C4 | 3.80 | — | Van der Waals |
| VAL 1527 | CA | U8 | C4 | 3.85 | — | Van der Waals |
| GLY 1231 | CA | G7 | C6 | 3.85 | — | Van der Waals |
| ASN 1233 | CG | G19 | C5 | 3.87 | — | Van der Waals |
| THR 1525 | CB | U8 | C6 | 3.89 | — | Van der Waals |
| MET 1232 | CE | G13 | C2' | 3.89 | — | Van der Waals |

Based on previous structures, two target recognition domain (TRD) loops bind the major groove of hemi-methylated DNA in the catalytic site of DNMT1. Here, in RNA-bound DNMT1, TRD loop 1 (1499-1514), which normally binds the phosphate backbone of DNA, is positioned toward the top lobe of the protein and away from the catalytic site in the RNA-bound DNMT1 (Extended Data Fig. 9a). Interestingly, there are two layers of extra density above the U quartet, the top layer of the pUG-fold. We were unable to model this density, but we hypothesize it may be comprised of other nucleotides from the (GU)_20_ RNA, as only 12 GU repeats are required to form the pUG-fold^22^. TRD loop 1 is not close enough to interact with the modeled pUG-fold but may be interacting with this unassigned density of hypothesized nucleotides (Extended Data Fig. 9b-c). TRD loop 2 (1528-1535), normally responsible for base-specific recognition in the catalytic site, is retracted away from the catalytic site as well (Extended Data Fig. 9d). The positioning of the TRD loops in the RNA-bound DNMT1 resembles that of the apo-, autoinhibited DNMT1 state (Extended Data Fig. 9e)^7^. Overall, the structural insights from the DNMT1 (Δ1-619)-pUG-fold RNA complex reveal an intricate set of interactions between the RNA and catalytically important residues of DNMT1 that render the active site incompetent for productive catalysis.

## Conformational dynamics of DNMT1 allow for promiscuous RNA binding

Motivated by the observation that mut3+CXXCmut DNMT1 adopts distinct RNA-bound and RNA-free classes, we hypothesized that the original full-length dataset would also contain closed and open states of DNMT1 bound to RNA. To capture these states, we reprocessed the data using a scheme like that used for the mut3+CXXCmut-(GU)_20_ dataset, with a specific focus on capturing RNA at the catalytic site (Supplemental Fig. 10). Indeed, we were able to see four different 3D classes: (1) closed conformation without RNA (closed-unbound), (2) closed conformation with RNA (closed-bound), (3) open conformation without RNA (open-unbound), and (4) open conformation with RNA (open-bound) (Fig. 5a-b). The closed-unbound state resembles the apo-, autoinhibited structures of DNMT1^7,14^ (Supplemental Fig. 11a) and our initial reconstruction, with the RNA density bound to the protein surface in the methyltransferase domain (Fig. 1b-c, Supplemental Fig. 11b-c). The RNA density in this map has the same overall shape and size as our previous reconstruction but is not as well-defined due to needing further classification. We did not classify these particles further to retain the prominence of the RNA density, as our objective in this reprocessing was to identify distinct conformational states.

Interestingly, the open-unbound state resembles the apo-, autoinhibited structures of DNMT1^7,14^ (Supplemental Fig. 11d-e). However, it is missing the top lobe of the RFTS domain, the CXXC domain, and the autoinhibitory linker, likely due to conformational flexibility. This indicates that the disengagement of the RFTS and CXXC domains, resulting in the open conformation of DNMT1, is not enough to position the catalytic loop and TRD loops in the catalytically competent configuration. This positioning is likely only achieved upon hemi- methylated DNA binding. The open-bound state resembles our maps of mut3+CXXCmut bound to (GU)_20_ RNA and the Δ1-619 DNMT1 bound to (GU)_20_ RNA (Supplemental Fig. 11f). The relative populations of these four states indicate that, even in the wild-type DNMT1-(GU)_20_ dataset, a detectable fraction of molecules spontaneously samples the open (RFTS/CXXC- disengaged) state, and a subset of these molecules bind (GU)_20_ RNA in the catalytic site.

To gain mechanistic insight into how mutations in DNMT1 mut3+CXXCmut alter the conformational dynamics underlying these states, we performed metadynamics molecular dynamics (MD) simulations, starting with the AlphaFold3-predicted structures of full-length wild-type DNMT1 and DNMT1 mut3+CXXCmut (Fig. 5c-d). In wild-type DNMT1, the conformational sampling profile showed a deep minimum at CV = 8.5 nm (state 2, Fig. 5c), corresponding to the closed (RFTS/CXXC-engaged), autoinhibitory state^31,32^. In contrast, the mut3+CXXCmut combination mutant minimum was shifted nearly 2 nm closer (CV = 10.3 nm) to the open, disengaged conformation (state 2 and 3, Fig. 5d), indicating that the mutations destabilize the autoinhibitory configuration and allow the enzyme to more readily sample the open, disengaged state. These observations support our cryo-EM structural analysis and explain the multiple modes of RNA binding to DNMT1, particularly the binding of RNA at the typically occluded catalytic site.

## DNMT1-interacting RNAs in cells have non-canonical G-quadruplex forming propensity

To determine whether DNMT1-bound RNAs in cells are predisposed to form G-quadruplexes and non-canonical G-quadruplex structures, such as pUG-folds, we analyzed published DNMT1 RNA immunoprecipitation sequencing (RIP-seq) data from human HL-60 cells ^16^, in which DNMT1-associated transcripts were partitioned into eight clusters by DNMT1 occupancy, promoter CpG methylation, and gene expression. We compared the two DNMT1- bound, highly expressed clusters: Cluster C (n = 2,737 genes; mean promoter methylation β = 0.21) and Cluster G (n = 584; β = 0.71) with the DNMT1-unbound Cluster A (n=600; β = 0.21). We then scored the DNMT1-bound RIP-seq peaks for the presence of GU or GA repeats of 11.5 or more (both are characteristic of a pUG-fold, but GA tracts have been shown to form pUG-folds with lower stability^23^) together with two independent machine-learning predictors – rG4detector, a CNN trained on in-cell DMS-MaPseq G-quadruplex data, and G4mer, an mRNAbert transformer trained on experimentally validated G-quadruplexes^33,34^ – as well as conventional rule-based G-quadruplex detectors such as cGcC and G4Hunter^35,36^. DNMT1-bound sequences were markedly enriched for predicted G-quadruplex structures or pUG-folds relative to methylation-matched DNMT1-unbound transcripts from cluster A. Within introns, 57% of Cluster C and 59% of Cluster G peaks scored positive with rG4 detector, compared with 34% of unbound pseudo-peaks (p < 0.001). G4mer showed a similar trend, with 34% of Cluster C and 37% of Cluster G peaks scoring positive, compared with 13% of unbound pseudo-peaks (Fig. 6a). To further confirm that DNMT1-bound peaks were enriched for G-quadruplex structures compared to sequences flanking the peaks, we generated a within-transcript comparison using flanking sequences 300-500 nucleotides (nt) away from the peaks which yielded similar results to our previous analysis (rG4detector: 57% versus 44% and 59% versus 46%; G4mer: 34% versus 19% for Cluster C, 37% versus 20% for Cluster G; all p < 0.001) (Extended Data Fig. 10a). We replicated these findings in an independent dataset, cell line, and assay, as reported by Wang et al., who identified DNMT1-interacting RNAs using enhanced cross-linking immunoprecipitation sequencing (eCLIP-seq) peaks^18^. In these DNMT1 eCLIP-seq peaks, 59% of intronic peaks scored positive by rG4 detector versus 26% of pseudo-peaks in genes without an eCLIP-seq peak, and 37% versus 11% by G4mer (p < 0.001; Fig. 6b). Rule-based scanners requiring canonical G-runs detected similar differences albeit at much lower sensitivity, indicating that the signal resides in non-canonical rather than classical G-quadruplexes (Supplemental Fig. 12)^35,36^.

**Fig. 5:**
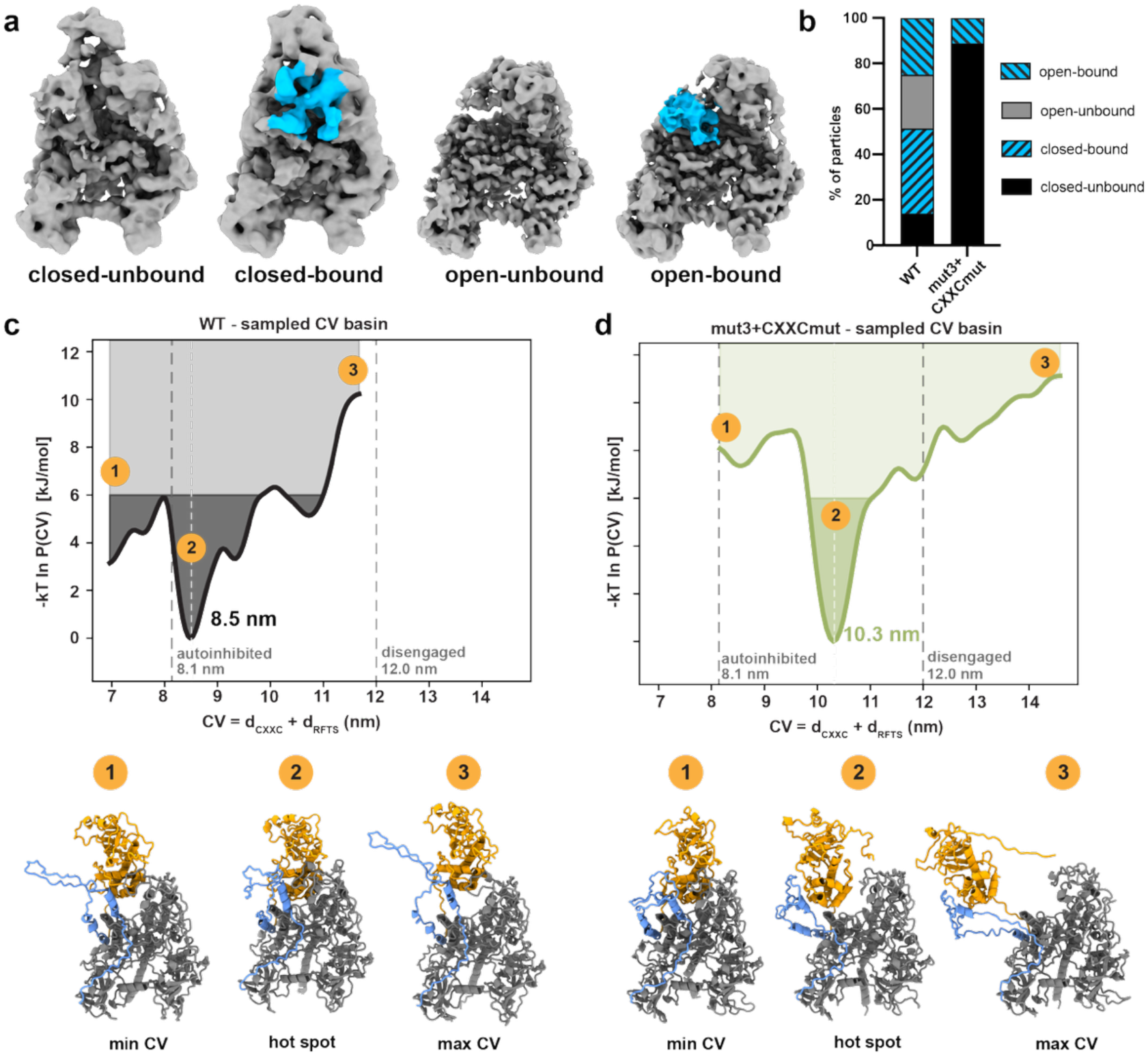
Conformational dynamics of DNMT1 can be observed in cryo-EM and metadynamics molecular dynamics. a, Maps of full length DNMT1 in the open or closed state, bound or unbound to pUG RNA. **b,** Quantification of particle fractions in each state from the wild-type DNMT1-(GU)_20_ and combination mutant mut3+CXXCmut DNMT1-(GU)_20_ RNA. **c,** Molecular dynamics energy landscape of wild-type DNMT1 (top) and conformational states at the indicated collective variable (CV) state (bottom). **d,** Molecular dynamics energy landscape of mut3+CXXCmut (top) and conformational states at the indicated CV state.

**Fig. 6:**
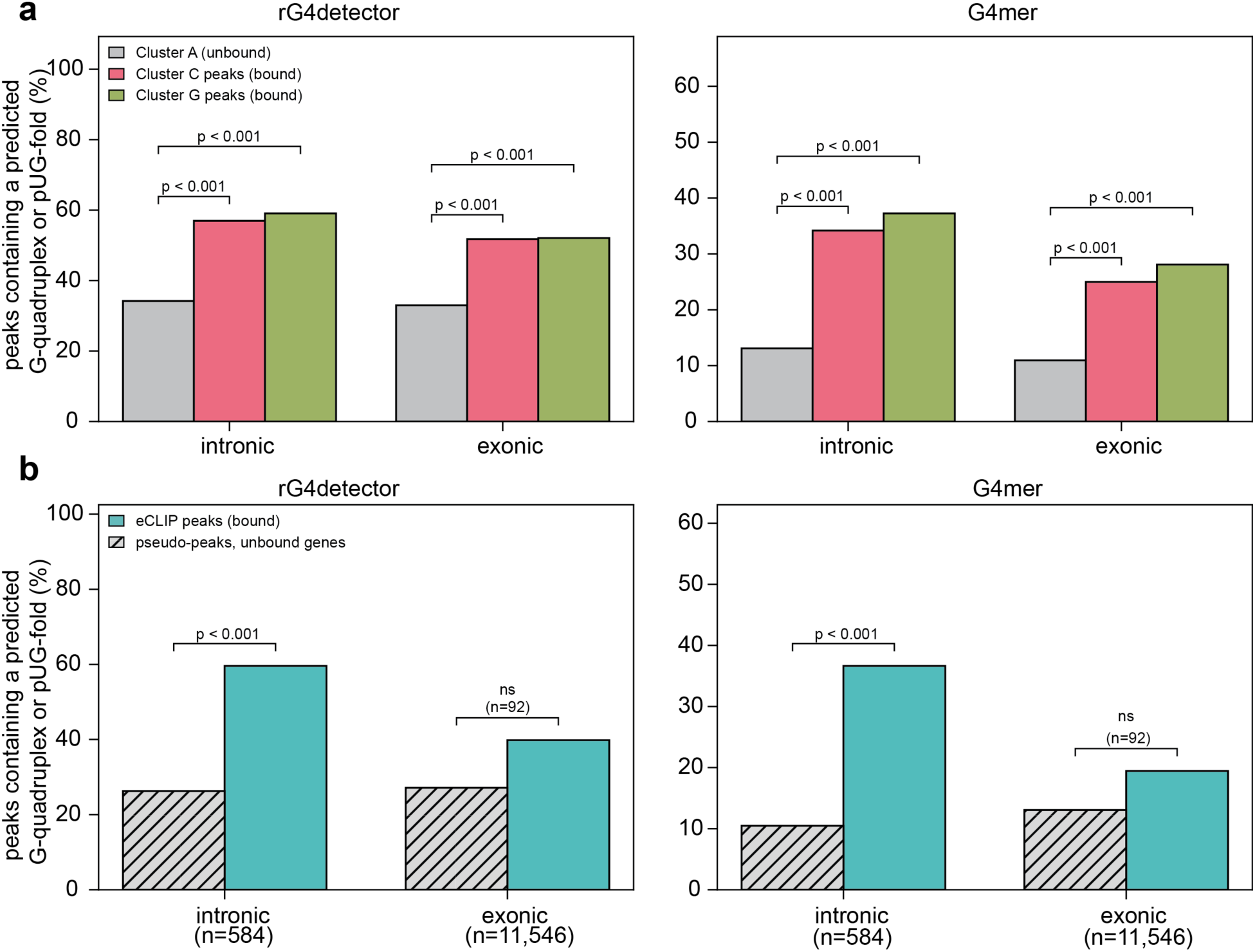
DNMT1-interacting RNAs in cells are enriched for G-quadruplex structures. **a,** Predicted percentage of canonical or non-canonical G-quadruplexes in RIP-seq peaks from HL-60 cells (Di Ruscio 2013) in cluster C and cluster G, versus pseudo-peaks placed at random positions in the pre-mRNA of cluster A genes. **b,** Predicted percentage of canonical or non-canonical G-quadruplexes in DNMT1 eCLIP peaks from HeLa cells (Wang et al. 2025) versus pseudo-peaks in genes carrying no eCLIP peak. Peak counts are given beneath each compartment.

To further validate these findings, we carried out a sliding-window analysis of the lncRNA ecCEBPA, which has been shown to regulate DNMT1^16^. ecCEBPA lncRNA (4786 nt, present in cluster C) is a known antagonist of DNMT1 activity at the *CEBPA* locus in cells and showed several regions throughout the lncRNA with a high propensity to form canonical or non-canonical G-quadruplex-like structures (Extended Data Fig. 10b-c). Interestingly, the previously identified R5 sequence in the 3’ extension, which binds DNMT1 with nanomolar affinity, is within one of the predicted G-quadruplex regions, hinting at the potential G-quadruplex-based mechanism of DNMT1 inhibition by this R5 region (Extended Data Fig. 10b-d). Together, these results establish that DNMT1-bound transcripts, both genome-wide and at individually characterized loci, have a propensity to form non-canonical G-quadruplex structures.

## Discussion

RNA has been identified as a regulator of DNMT1 *in vivo*, despite the protein lacking a well-defined RNA-binding domain. Structured RNAs have been proposed to bind to the catalytic domain, but no studies have examined these interactions through a structural lens^16,19^. Additionally, the structural and mechanistic basis of RNA-DNMT1 interactions that lead to inhibition of methyltransferase activity has remained unknown.

Here, we examined the inhibitory, non-canonical G-quadruplex pUG-fold RNA binding to DNMT1 with cryo-EM. We found that pUG-fold RNA can bind to DNMT1 in two ways: (1) on the protein surface and (2) in the catalytic site (Fig. 1,3-4). When DNMT1 is in the closed, autoinhibited conformation, the RNA can dynamically bind across the positively charged belt between the CXXC and methyltransferase domains, thereby blocking DNA binding. When the RFTS and CXXC domains are disengaged from the methyltransferase domain, thereby rendering the protein in an open state, the RNA can nestle into the catalytic site and compete with DNA for binding (Fig. 7a). Therefore, the location of RNA binding is dependent on the conformational dynamics of the protein, but both locations result in DNMT1 inhibition. Although we initially thought that the structure of the DNMT1-RNA complex would guide the design of a separation-of-function mutant to further test the importance of RNA binding in cells, the multiple RNA binding sites on DNMT1 prevented our repeated attempts to disrupt RNA binding without disrupting DNA methyltransferase activity.

**Fig. 7:**
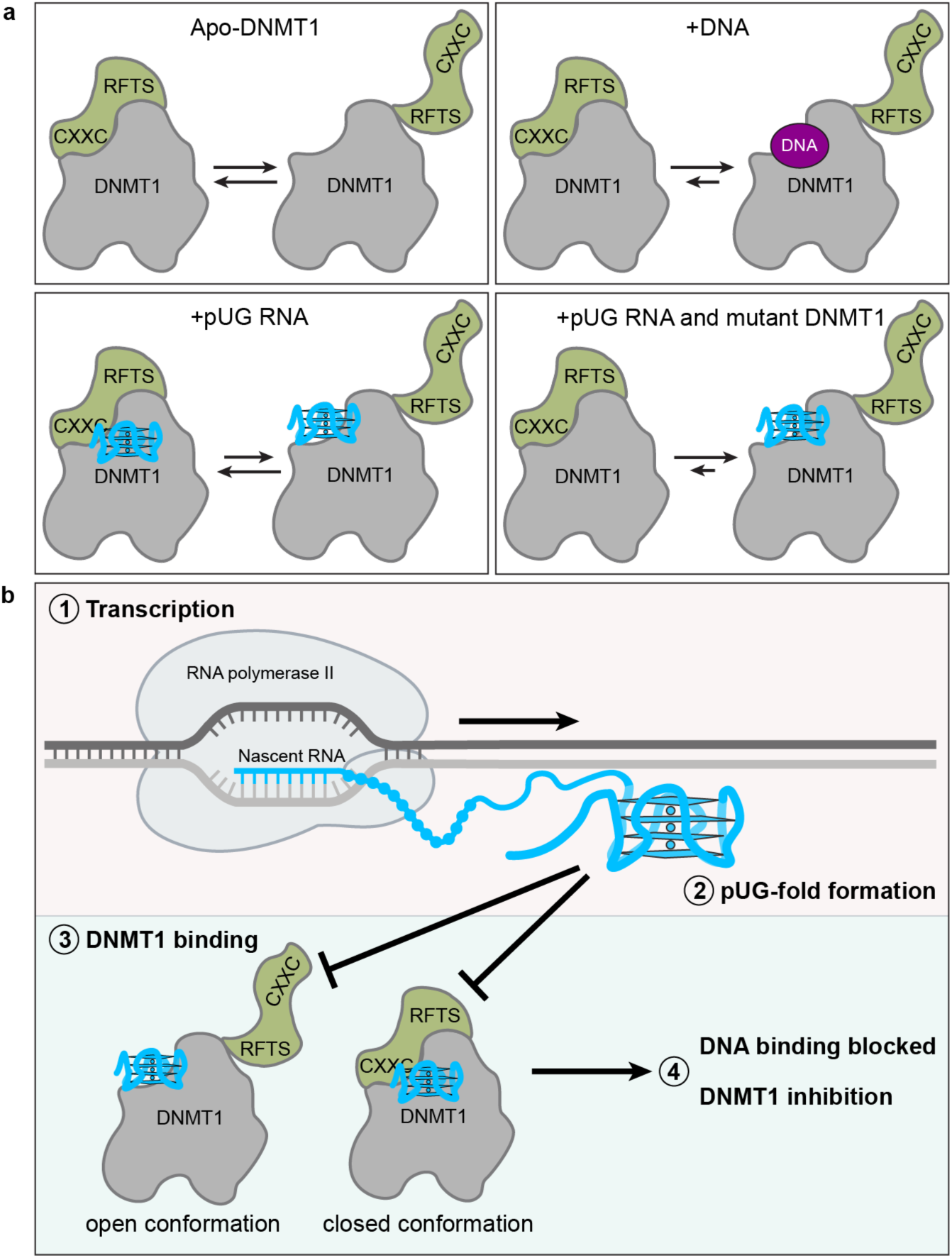
Conformational dynamics of DNMT1 allow for promiscuous RNA binding. a, In the apo- state, DNMT1 samples both the open and closed conformations. DNA binding locks DNMT1 into the open conformation. pUG-fold RNA binds DNMT1 in both open and closed conformations. The mut3+CXXCmut mutations inhibit RNA binding at the mutated, closed site, so RNA binds DNMT1 in the open conformation. **b,** Proposed model for biological significance. (1) RNA polymerase II transcribes a gene and (2) the nascent mRNA folds into a pUG-fold RNA. (3) The pUG-fold binds to DNMT1 in the open or closed conformation to (4) block DNA binding and result in DNMT1 inhibition.

In this study, we utilized AlphaFold3 to guide cryo-EM experiments and structure determination. AlphaFold3 predicted two RNA-binding sites in the full-length DNMT1 construct, which matched the results from 3D variability analysis showing the binding of pUG-fold RNA between the CXXC and methyltransferase domains (Fig. 2b-c). We utilized these predictions to guide mutagenesis to disrupt RNA binding. AlphaFold3 predictions for the combination mutant with TERRA RNA indicated that the mutated residues no longer interact with RNA; however, the RNA was still predicted to be positioned near the methyltransferase domain, albeit with lower confidence (Extended Fig. 5d, Supplemental Fig. 5). Interestingly, when using a Δ1-619 truncation mutant or truncation mutant Δ1-619 containing the mut3+CXXCmut mutations, AlphaFold3 predicted the RNA would bind in the catalytic site (Extended Fig. 5d, Extended Fig. 7b, Supplemental Fig. 6, 8-9), which motivated us to pursue cryo-EM studies of the double mutant and Δ1-619 DNMT1 in complex with pUG-fold RNA. Overall, this study demonstrates the utility of AlphaFold3 in guiding cryo-EM experiments.

We resolved a 2.5 Å map of pUG-fold RNA bound to the catalytic site of DNMT1, which allowed us to propose a mechanism for the inhibition of DNMT1 by pUG-fold RNA (Fig. 4). We identified many residues that interact with the RNA, several of which are also implicated in DNA binding (Fig. 4e-g, 4j-k, Extended Data Fig. 8, Table 3). This indicates that pUG-fold RNA inhibits DNMT1 by competing with the DNA for binding in the catalytic site and engaging with residues critical for DNA coordination, providing a structural explanation for the competitive inhibition seen biochemically ^21^. This binding mode only occurs when the RFTS and CXXC domains disengage from the catalytic domain and open the catalytic site for binding. Disengagement of the RFTS and CXXC domains is likely insufficient to render DNMT1 catalytically competent, and the binding of pUG-fold RNA stabilizes and reinforces this catalytically incompetent state. This mechanism of inhibition differs from that of current DNMT1 small-molecule inhibitors. Nucleoside inhibitors, such as 5-azacytidine and decitabine, incorporate into the DNA and form covalent bonds with DNMT1 to inhibit methyltransferase activity^37^. The non-nucleoside inhibitor GSK-3484862 competes with the catalytic loop to intercalate into the DNA, while also interacting with the TRD loops^30^. While these inhibitors require DNA binding for inhibition, pUG-fold RNA-mediated inhibition of DNMT1 acts in a DNA-free manner. Perhaps the design of future small-molecule inhibitors of DNMT1 can be guided by the interactions between DNMT1 and the pUG-fold.

We presented the first cryo-EM reconstruction of a non-canonical RNA G-quadruplex and of pUG-fold RNA recognition by a protein. The catalytic site of DNMT1 forms a cleft of loops around the RNA that interact with the phosphodiester backbone, looped-out uracil nucleotides, and guanine tetrads. More specifically, R1269 binds the phosphate groups from two different G tetrad layers, creating the dominant interaction between DNMT1 and the RNA (Extended Data Fig. 8a-b). This residue has been previously implicated in binding phosphate groups in the context of hemi-methylated DNA^25^. Additionally, the aromatic edge of F1274 stacks against the G7 base of the G1 tetrad to form a T-shaped aromatic interaction, which may confer DNMT1 specificity in binding G-quadruplex RNAs beyond the pUG-fold (Extended Data Fig. 8a). Future studies may determine if DNMT1 binds to canonical G-quadruplexes in a similar manner and examine why DNMT1 is more strongly inhibited by pUG-fold RNA compared to canonical G-quadruplexes.

Interactions of other proteins with DNA and RNA G-quadruplexes are more commonly either structure- or sequence-specific. Many structures of thrombin bound to DNA aptamers that form G-quadruplexes show dominant interactions between the protein and the quadruplex lateral loops. A cryo-EM structure of nucleolin with the G-quadruplex-forming Myc DNA found that nucleolin binds the lateral loops and the phosphate backbone of the G-quadruplex^38^. In contrast, the RNA helicase DHX36 uses an α-helical element, the DHX36-specific motif (DSM), that packs against the 5′ G-tetrad face of both RNA and DNA G-quadruplexes, a mode of G-quadruplex structure recognition that is proposed to be more general. This tetrad-face contact works with two additional groove-binding loops, OI and RecA2-G4, to further engage G- quadruplexes^39^. Likewise, superfamily 1 (SF1) helicase Pif1 recognizes all three G tetrads of a G-quadruplex for general structure recognition^40^. It will be intriguing to investigate whether other pUG-fold binding proteins, like DNMT1, recognize the sequence-specific flipped-out uracil bases, as well as the overall structure of the pUG-fold.

pUG-fold RNAs have been previously identified as a driver of gene silencing in *C. elegans* by recruiting RNA-dependent RNA polymerase (RdRP) which synthesizes small interfering RNAs (siRNAs)^22,41^. While there is no RdRP ortholog in humans, a previous study identified more than 20,000 pUG-fold sequences in mammalian introns^22^. This number may be an underestimate, given that pUG-folds can tolerate deviations in sequence^23^. It has not yet been shown that GU repeats exist as pUG-fold structures in living cells; however, this may be due to dynamic folding and unwinding by helicases. This is similar to canonical G-quadruplexes, as studies have found that some G-quadruplexes are folded in cells while others are globally unfolded^42–44^.

We utilized published RIP-seq and eCLIP-seq data to characterize the nature of DNMT1-RNA interactions in cells. Machine-learning predictors, rG4detector and G4mer, show that the RNA sequences DNMT1 contacts in cells are enriched for canonical and non-canonical G-quadruplexes (Fig. 6). We found that the DNMT1 antagonist lncRNA ecCEBPA contains several regions with a propensity to form G-quadruplexes (Extended Fig. 10). Future studies should examine whether ecCEBPA forms G-quadruplexes and whether ecCEBPA-mediated DNMT1 inhibition at the *CEBPA* locus depends on G-quadruplex formation (Fig. 7b).

In summary, our work has determined a structural basis for pUG-fold RNA-mediated regulation of DNMT1 across different DNMT1 conformations. This is the first cryo-EM -We also show that previously identified DNMT1-interacting RNAs in cells have a propensity to form canonical and non-canonical G-quadruplexes. Overall, these findings position DNMT1 alongside PRC2, LSD1, and other chromatin modifiers as a direct target of ncRNA-mediated regulation, integrating the DNA methylation machinery into the broader landscape of RNA-guided chromatin control that governs genome function (Fig. 7b)^45–48^.

## Data availability

Cryo-EM maps and fitted models were deposited to the EM Data Bank (under accession numbers EMD-78737, EMD-78766, EMD-78750) and the PDB (under accession numbers 38DB, 38EM, 38DT). Corresponding accession codes for each structure can be found in Table 1.

## Acknowledgements

We thank A. Herbst of the University of Colorado Boulder Biochemistry Shared Instrument Pool Facility (RRID: R24OD033699-01) for training and access to shared instrument facilities and T. Nahreini of the University of Colorado Boulder Biochemistry Cell Culture Facility (RRID: SCR_018988). We also thank E. Hartwick and S. Laursen of the University of Colorado Boulder Biochemistry Krios Electron Microscopy Facility (RRID: SCR_019057) for cryo-EM data screening and maintenance of computational infrastructure, as well as G. Morgan and S. Zimmerman of the University of Colorado Boulder Electron Microscopy Services. Cryo-electron microscopy data were collected at the Pacific Northwest Cryo-EM Center (PNCC), supported by NIH grant U24GM129547 and located at the Environmental Molecular Sciences Laboratory (EMSL), a DOE Office of Science User Facility sponsored by the Office of Biological and Environmental Research. We thank the PNCC staff for assistance with microscope operation and data collection. We also thank members of the Kasinath and Cech labs, especially Ella Tommer, for their assistance in experiments.

## Author contributions

J.S., T.R.C., and V.K. conceived the study and wrote the manuscript. J.S, and V.K analyzed the data. J.S performed all the biochemical and cryo-EM structural experiments. J.S and V.K performed the MD simulations and computational analysis. J.S and V.K wrote the manuscript. J.S, T.R.C, and V.K read and edited the manuscript.

## Materials and Methods

### Protein expression and purification

Full-length human DNMT1 (UniProt P26358-1), truncation, and mutant constructs were cloned into a pFastBac vector (Invitrogen Life Technologies) with N-terminal 6xHis, maltose binding protein (MBP) tags to produce recombinant baculovirus. Baculovirus stocks were generated in Sf9 cells and used to infect 1 L of Trichoplusia ni (HighFive) cells at a density of 1 million cells/mL and incubated for 66 hours at 27°C, 120 revolutions per minute (rpm). Cells were harvested by centrifugation for 30 minutes at 3000 rpm at 4°C, washed with cold buffer (25 mM HEPES pH 7.5, 150 mM NaCl), frozen in liquid nitrogen and stored at -80°C until use.

All purification steps were done at 4°C. Cell pellets were resuspended in cold lysis buffer (25 mM HEPES pH 7.5, 250 mM NaCl, 1 mM TCEP, 0.5% NP-40, 10% glycerol, supplemented with cOmplete Mini, EDTA-free protease inhibitor cocktail tablet (Roche), 0.2 mM PMSF, 10 µM leupeptin, 1 µM aprotinin, 1 mM pepstatin, and DNase) and incubated for 1 hour with rotation. Cells were sonicated four times at 60% power with 10 sec ON and 20 sec OFF (Q-Sonica). Cell lysate was cleared by centrifugation at 15,000 rpm for 30 min. The pellet was discarded and the supernatant was incubated with equilibrated amylose resin (NEB, E8021L) for one hour with rotation. The resin was washed with 5 column volumes (CV) of low salt buffer (25 mM HEPES pH 7.5, 150 mM NaCl, 1 mM TCEP, 10% glycerol, supplemented with protease inhibitors) with 0.01% NP-40, 5 CV of high salt buffer (25 mM HEPES pH 7.5, 1 M NaCl, 1 mM TCEP, 10% glycerol, supplemented with protease inhibitors), and 5 CV of low salt buffer. Protein was eluted with 5 CV of elution buffer (25 mM HEPES pH 7.5, 150 mM NaCl, 1 mM TCEP, 10% glycerol, 10 mM maltose). To remove the 6xHis and MBP tags, eluted protein was incubated with 3C protease (Cytiva) with 1:100 w/w ratio ON with rotation. Cleaved protein was purified by a HiTrap 5 mL Heparin HP column (Cytiva) and fractions were pooled and concentrated. The concentrated fractions were injected into a Superose 6 Increase 10/300 column equilibrated with 25 mM HEPES pH 7.5, 150 mM NaCl, 1 mM TCEP, 10% glycerol). Peak fractions were pooled, concentrated, aliquoted and stored at -80°C until further use.

## RNP complex assembly

For the full-length DNMT1-RNA structure, (GU)_20_ -A_20_-3’ Biotinylated RNA was purchased from IDT: ([5’ GUGUGUGUGUGUGUGUGUGUGUGUGUGUGUGUGUGUGUGUAAAAAAAAAAAAAAAAAAAA-Biotin 3’]). (GU)_20_ RNA was used to increase the number of particles bound to RNA, due to DNMT1 having a higher affinity for (GU)_20_ versus (GU)_12_ RNA^21^. RNA was resuspended in water and heated at 95°C for 5 min, snap cooled on ice for 3 min, refolded in buffer (25 mM Hepes pH 7.5, 100 mM KCl, 2.5 mM MgCl_2_) and incubated at 37°C for 30 min. RNA, DNMT1, and S-adenosyl methionine homocysteine (SAH) were added together for a final concentration of 2 µM, 1.7 µM, and 40 µM , respectively. The binding reaction was incubated at room temperature for 30 min. 25% glutaraldehyde was diluted to 2.5% with 25 mM HEPES, added to the binding reaction for a final concentration of 0.1%, and incubated on ice for 10 minutes. The reaction was quenched with 25 mM Tris pH 8.

For the mut3+CXXCmut DNMT1-RNA structure, (GU)_20_ RNA was purchased from IDT (5’GUGUGUGUGUGUGUGUGUGUGUGUGUGUGUGUGUGUGUGU 3’). RNA was folded as stated above. RNA, mutant DNMT1, and S-adenosyl methionine homocysteine (SAH) were added together to give a final concentration of 2.5 µM, 2 µM, and 80 µM, respectively. The binding reaction was incubated at room temperature for 30 min.

For the Δ1-619 DNMT1-RNA structure, (GU)_20_ RNA was folded as stated above. RNA, Δ1-619 DNMT1, and S-adenosyl methionine homocysteine (SAH) were added together for a final concentration of 9.8 µM, 5 µM, and 80 µM, respectively. The binding reaction was incubated at room temperature for 30 min.

## Cryo-EM sample preparation

For the full-length DNMT1-RNA structure, Quantifoil 1.2/1.3 Au 300 mesh grids were cleaned with chloroform and ethanol. Grids were glow discharged used Tergeo EM plasma cleaner. 4 µL of the crosslinked binding reaction was applied to the grid. The grid was incubated for 30 sec in a Vitrobot Mark IV humidified chamber, blotted for 4-5 s at 4°C and 100% humidity, and then plunged into liquid ethane.

For the mut3+CXXCmut DNMT1-RNA and Δ1-619 DNMT1-RNA structures, Quantifoil 1.2/1.3 Au 300 mesh grids were prepared as stated above. 4 µL of the reaction was applied to the grid. The grid was incubated for 30 sec in a Vitrobot Mark IV humidified chamber, blotted for 2-3 s at 4°C and 100% humidity, and then plunged into liquid ethane.

## Cryo-EM data collection and processing

All cryo-EM datasets were collected at the Pacific Northwest Cryo-EM Center (PNCC) on a Titan Krios equipped with Gatan K3 direct detector and BioContinuum energy filter. All datasets used a 10 eV filter slit width. Movies were recorded in super-resolution mode at a nominal magnification of 130,000x. The full-length DNMT1-RNA dataset was collected at a calibrated pixel size of 0.6488 Å/pixel (super resolution 0.3244 Å/pixel). The Δ1-619 DNMT1-RNA dataset was collected at a calibrated pixel size of 0.8257 Å/pixel (super resolution 0.41285 Å/pixel). The double mutant DNMT1-RNA dataset was collected at a calibrated pixel size of 0.6485 Å/pixel (super resolution 0.32425 Å/pixel). Automated data collection was done using SerialEM with a defocus range of -0.6 to -2.0 µm for the full-length DNMT1-RNA dataset and -0.8 to -2.5 µm for both the Δ1-619 DNMT1-RNA and double mutant DNMT1-RNA dataset. A total dose of 50 electrons per square angstrom (e-/Å^2^). Data were collected as dark-subtracted, non-gain corrected compressed movies (.tif format). All movies were motion corrected and CTF corrected in CryoSparc v4.7^49^. Further processing steps were done in CryoSparc v4.7.

For the full-length DNMT1-RNA reconstruction, only micrographs with a 4 Å or better CTF fit and a maximum relative ice thickness of 1.2 were retained for further analysis. Particles were picked from all micrographs and went through one round of 2D classification to remove junk particles. A subset of particles was used to generate ab initio reconstructions that were used for two rounds of heterogeneous refinement. Multiple rounds of 3D variability analysis were performed to investigate continuous flexibility in the dataset. Clusters showing an extra density in the catalytic domain were analyzed further. Particles were extracted without binning and then underwent nonuniform refinement and local filtering.

For the double mutant-RNA reconstruction, only micrographs with an 8 Å CTF fit resolution, a maximum relative ice thickness of 1.535, and intrinsic ice defect area less than 30% were retained for further analysis. Particles were picked from a subset of micrographs to generate 2D templates to pick from the entire dataset. One round of 2D classification, ab initio reconstruction, and heterogeneous refinement were done to isolate good particles before extracting particles without binning. Subsequent processing steps include global CTF refinement and nonuniform refinement. Particles were classified into ten classes without alignment and grouped into two conformations: an RFTS/CXXC-disengaged state bound to RNA and an RFTS/CXXC-engaged state with no RNA bound. Each conformation was refined further through non-uniform refinement and local filtering.

For the Δ1-619 DNMT1-RNA reconstruction, only micrographs with a 7 Å or better CTF fit and a maximum relative ice thickness of 1.2 were retained for further analysis. Particles were picked from a subset of micrographs to generate 2D templates to pick from the entire dataset. Two rounds of 2D classification, ab initio reconstruction, and heterogeneous refinement were done to isolate good particles before extracting particles without binning. Subsequent processing steps include global CTF refinement and nonuniform refinement. Particles were subset into 2 groups by per-particle scale and refined separately. One subset that yielded a better map was used for further processing, including focused 3D classification on the RNA density, non-uniform refinement, and local filtering.

## Model building

ModelAngelo was used to generate an initial model for the full-length DNMT1 from the final consensus refinement map from CryoSparc^50^.All the coordinates were adjusted and rebuilt using COOT v1.1.20. The coordinates of the pUG-fold RNA (PDB: 8TNS) were rigid body docked into the cryo-EM map in ChimeraX^51^ and merged with the full-length DNMT1 model. The merged coordinates were refined with Phenix Real Space Refinement^52^.

The Δ1-619 DNMT1-RNA model was built using an AlphaFold3 prediction of apo-Δ1-619 DNMT1. All coordinates were adjusted and rebuilt using COOT v1.1.20^53^. The coordinates of the pUG-fold RNA (PDB: 8TNS) were flexibly fit using ISOLDE^54^. These coordinates were merged with the Δ1-619 DNMT1 protein coordinates in COOT. The merged coordinates were refined with Phenix Real Space Refinement. This model was subsequently used for modeling mut3+CXXCmut DNMT1-RNA, adjusted in COOT, and refined with Phenix Real Space Refinement.

## Mass photometry

Samples were diluted to a final concentration of 1 nM in freshly made 25 mM HEPES pH 7.5, 150 mM NaCl, 1 mM TCEP buffer for mass photometry. Glass coverslips (24 × 50 mm, Thorlabs Inc. 71861-054) were washed 3 times with Milli-Q water and HPLC-grade isopropanol, then dried with a clean stream of compressed air. Sample chambers were assembled by placing clean 6-well silicon gaskets (Fisher Scientific, NC2754003) on the cleaned coverslip, with the gasket positioned on the stage of the mass photometer. Measurements were performed using a TwoMP mass photometer (Refeyn LTD, Oxford, UK). Data were acquired using the Acquire MP software package and analyzed using the Discover MP software, both from Refeyn. 20 µL of the sample was applied, and the mass photometer was focused. Data acquisition was started immediately to record a 60-second movie. The mass calibration was achieved using beta-amylase (Sigma-Aldrich, A3176, 10 nM in buffer; 56, 112, and 224 kDa). All measurements were performed at room temperature.

## Electrophoretic mobility shift assay (EMSA)

DNA containing a single hemi-methylated site (1 5mC DNA) was purchased from IDT as a duplex ([5’ TACGTATCCGTATCCGGTTA/5mC/GTATCCGAATCCGTACCGT 3’] [5’ACGGTACGGATTCGGATACGTAACCGGATACGGATACGTA]). To end label oligos, 50 pmol of either (GU)_20_ RNA or 1 5mC DNA was incubated for 30 minutes at 37°C with 10 U of T4 PNK (NEB) and 1 µL of γ-^32^P-ATP in 20 µL1× PNK buffer. Radiolabeled oligos were purified using Micro Bio-Spin 6 columns (BioRad). Radioactivity was measured by a scintillation counter.

End-labeled RNA was heated at 95°C for 5 min, snap cooled on ice for 3 min, and refolded in 1x refold buffer (50 mM Tris-HCl pH 7.5, 100 mM KCl, 2.5 mM MgCl_2_, 0.1 mM ZnCl_2_, 2.0 mM BME, 0.05% NP40, 5% v/v glycerol) at 37°C for 30 min. Each binding reaction was set up with the indicated amount of DNMT1 and 1000 cpm of end-labeled RNA or DNA in binding buffer (50 mM Tris-HCl pH 7.5, 100 mM KCl, 2.5 mM MgCl_2_, 0.1 mM ZnCl_2_, 2.0 mM BME, 0.05% NP40, 5% v/v glycerol, 0.1 mg/mL BSA [NEB], 0.1 mg/mL baker’s yeast tRNA [Sigma R5636]). Reactions were incubated for 30 minutes at 30°C. Binding reactions were run on a 1% agarose gel in 1x TBE for 90 minutes at 66 V at 4°C. The gel was dried, exposed overnight, and imaged on an Amersham Typhoon. All EMSAs were performed in triplicate and quantified with GelAnalyzer-23.1.1. Binding curves were calculated using Prism 10.

## Binding competition assay

For testing (GU)_20_ RNA competition of protein pre-bound to hemi-methylated DNA (sequence above), (GU)_20_ RNA was heated at 95°C for 5 min, snap cooled on ice for 3 min, refolded in 1x refold buffer (above) at 37°C for 30 min. Each binding reaction initially contained 500 nM WT DNMT1 or 1000 nM mut3+CXXCmut DNMT1 and 1000 cpm of end-labeled DNA, incubated at room temperature for 30 minutes. After 30 minutes, the indicated concentration of RNA in 1X binding buffer (above) was added and incubated at room temperature for one hour.

The same initial steps of RNA folding were repeated to test hemi-methylated DNA competition of protein pre-bound to (GU)_20_ RNA. Each binding reaction contained 125 nM WT DNMT1 or mut3+CXXCmut DNMT1 and 1000 cpm of end-labeled RNA, incubated at room temperature for 30 minutes. After 30 minutes, the indicated concentration of hemi-methylated DNA in 1X binding buffer was added and incubated at room temperature for one hour.

Binding reactions were run on a 1% agarose gel in 1x TBE for 90 minutes at 66 V at 4°C. The gel was dried, exposed overnight, and imaged on an Amersham Typhoon. All EMSAs were performed in triplicate and quantified with GelAnalyzer-23.1.1. Binding curves were calculated using Prism 10.

## Molecular Dynamics Simulations

Initial structures for full-length DNMT1 wild-type and mut3+CXXCmut were prepared from AlphaFold3 prediction. Structures were protonated at pH 7.5 using PROPKA3 and parameterized with the AMBER ff19SB force field and the OPC explicit water model using tLeap (AMBER 22). Each system was solvated in a truncated octahedron box with a 10 Å buffer, neutralized with Cl⁻ ions, and supplemented with 192 Na⁺/Cl⁻ pairs (∼150 mM ionic strength). Energy minimization was performed in two stages (restrained then unrestrained), followed by gradual heating from 0 to 300 K over 100 ps under NVT conditions with harmonic restraints on heavy atoms. Five sequential NPT equilibration stages (1 ns each) progressively released positional restraints. Production MD was performed using pmemd.cuda under NPT conditions (300 K, 1 bar) with a 2 fs timestep, SHAKE constraints on hydrogen bonds, PME electrostatics, and a 9 Å nonbonded cutoff. Each system was simulated for a minimum of 120 ns prior to metadynamics simulations.

## Well-Tempered Metadynamics

Metadynamics simulations were performed using OpenMM 8.2.0 and OpenMM-Plumed 2.1 with the PLUMED 2.9.2 plugin. To maximize computational efficiency, explicit solvent and counterions were stripped and simulations were conducted in the NVT ensemble at 300 K using a Langevin integrator (2 fs timestep) with the protein-only topology (∼25,450 atoms). Coordinates were initialized from the final frame of production MD.

The collective variable (CV) was defined as the sum of two Cα center-of-mass distances: CV = d(CXXC, MTase core) + d(RFTS, MTase core), with CXXC (aa 621–700), RFTS (aa 352–620), and MTase core (aa 1100–1500) serving as the domain groups. The MTase core (401 Cα atoms) served as a stable positional anchor. The autoinhibited baseline was CV ≈ 8.1 nm; CXXC/RFTS disengagement was defined as CV ≥ 12.0 nm.

Gaussian hills were deposited every 500 integration steps (1 ps) with a width of σ = 0.25 nm and a bias factor of γ = 15. A harmonic lower-wall restraint was applied to prevent artificial compaction. System-specific parameters were as follows:

DNMT1 mut3+CXXCmut:

Initial hill height H = 2.4 kJ/mol; lower wall at CV = 7.0 nm (κ = 2000 kJ/mol/nm²); achieving full disengagement (CV = 12.9 nm) within 23 ns.

DNMT1 WT: H = 2.4 kJ/mol, lower wall at CV = 7.0 nm (κ = 2000 kJ/mol/nm²), with no disengagement observed over 150 ns of cumulative sampling.

## DNMT1-bound transcript clusters

Gene clusters were taken from Di Ruscio *et al.*, DNMT1 RIP-seq in HL-60 cells (GEO: GSE321262) and were defined by the published intersection of DNMT1 RIP-seq occupancy, promoter CpG methylation (RRBS, mean β), and transcript expression^16^. No re-clustering was performed. Analysis was restricted to two DNMT1-bound clusters: C (bound, hypomethylated, high expression; *n* = 2,737 genes), G (bound, hypermethylated, high expression; *n* = 584), and the DNMT1-unbound hypomethylation cluster A (unbound, hypomethylated). Other unbound clusters (E, F) and the high-methylation/low-expression background cluster B were excluded. Clusters C, G versus A form the primary comparison.

## RNA sequence windows scored

G-quadruplex propensity was scored on DNMT1-bound pre-mRNA rather than spliced transcripts, because DNMT1 RIP-seq peaks are predominantly intronic (51.7% intronic, 8.1% exonic, 40.2% intergenic). Peak coordinates (137,854 peaks, HL-60, hg19) were from Di Ruscio et al. 2013 (GSE321262); the assembly was confirmed by exon-overlap enrichment (2.86-fold hg19 versus hg38)^16^. Gene symbols were mapped to hg19 refGene loci, and genomic spans, including introns, were retrieved and reverse-complemented for minus-strand genes.

## G-quadruplex and non-canonical G-quadruplex propensity scoring

Four separate methods were applied to identical input sequences: two machine-learning predictors and two rule-based motif scanners, the latter used as specificity controls for canonical G-run architecture.

Each peak was tiled into 150-nt windows at 10-nt steps, the fixed input length of both predictors, and scored with rG4detector (CNN trained on rG4-seq; threshold 1.3) and G4mer (transformer; threshold 0.5). A peak was classified as containing a predicted G-quadruplex or pUG-fold if any window exceeded threshold, or if the peak body contained a GU or GA tract of at least 11.5 dinucleotide units. Peaks with multiple qualifying windows or tracts count once, so percentages are the fraction of peaks that score positive. Rule-based scanners cGcC (threshold 4.5) and G4Hunter (0.9) were applied to identical windows for canonical-architecture comparison; cGcC is undefined without cytosine, and those windows were excluded.

## Control sequence sets

Each of the three controls answered a distinct question. Unbound-gene control: 600 cluster A genes, unbound but matched to cluster C on promoter methylation (beta 0.198 versus 0.207, p = 0.62), carrying pseudo-peaks whose counts and widths were drawn from the empirical peak distribution and placed at random positions. Within-transcript control: flanking regions 300-500 nt from the nearest peak center, overlapping no peak anywhere. Independent replication: DNMT1 eCLIP-seq peaks from HeLa cells (GEO: GSE169102; 13,023 peaks, hg38, Wang et al.; assembly verified by exon fraction 0.916 hg38 versus 0.124 hg19) versus pseudo-peaks in 250 genes with no eCLIP-seq peak, span-matched to the bound distribution (p = 0.92)^18^.

Statistical comparisons used two-sided Fisher exact tests on peak counts, testing the plotted quantity directly. Windows spanning an exon-intron boundary (exon fraction between 0 and 0.5) were counted but excluded from compartment comparisons.

## Software

Analyses used Python 3 with NumPy, pandas, SciPy (mannwhitneyu, fisher_exact,hypergeom) and matplotlib; bedtools for interval operations. Processing scripts will be made available at the Kasinath Lab GitHub (https://github.com/KasinathLab/DNMT1-interacting-RNAs-G4analysis).

---

**Extended Data Fig. 1:**
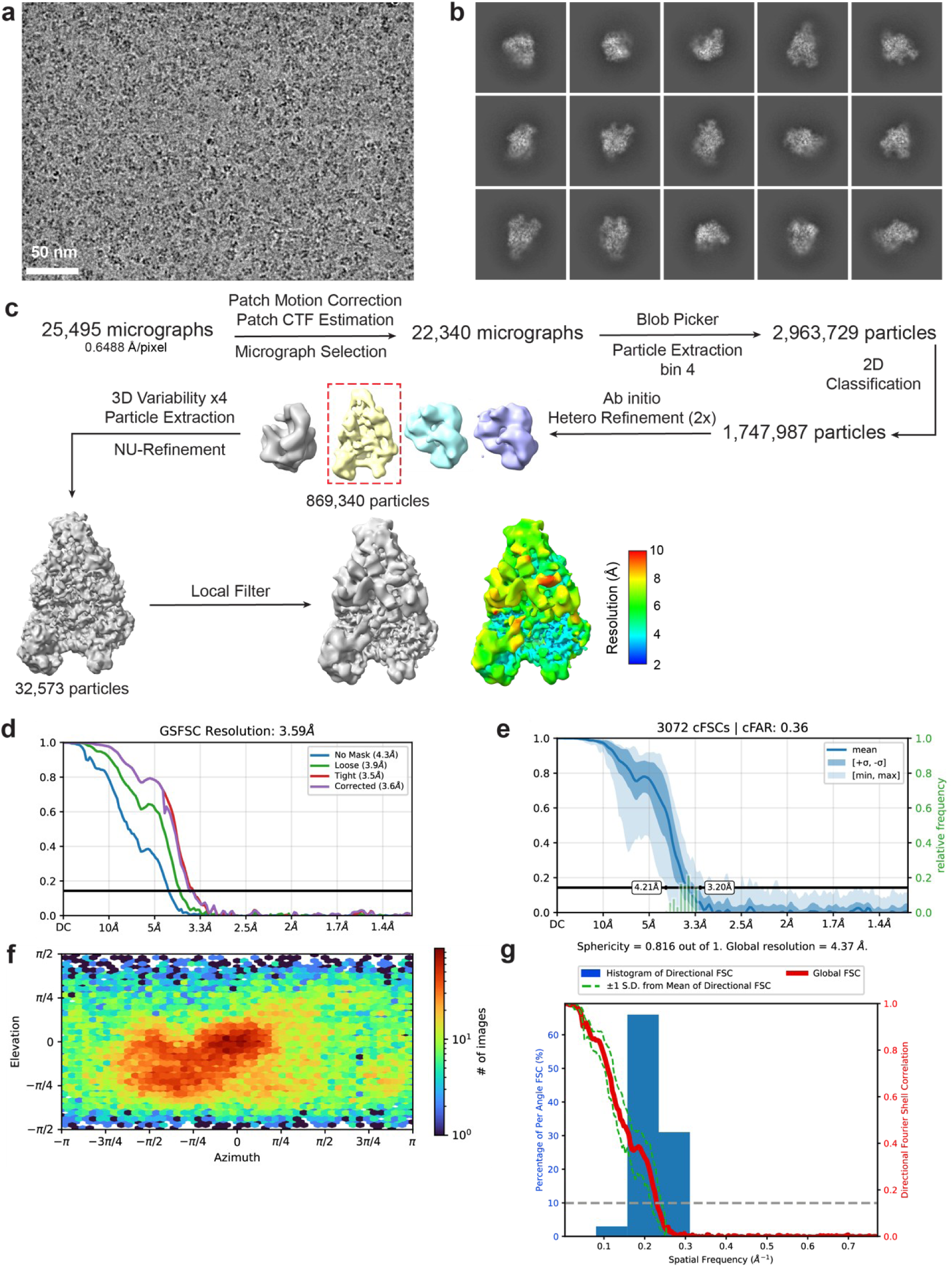
Data processing scheme for full-length DNMT1 and pUG-fold RNA. a, Representative micrograph. **b,** Representative 2D projections. **c,** Data processing scheme. Local filtered maps are colored by resolution. All data processing steps were done in CryoSPARC v4.7.1. **d,** Fourier shell correlation (FSC) plot. **e,** Conical Fourier shell correlation (cFSC) plot. **f,** Posterior precision directional distribution plot. **g,** 3D FSC for the final NU- refinement map obtained from CryoSPARC.

**Extended Data Fig. 2:**
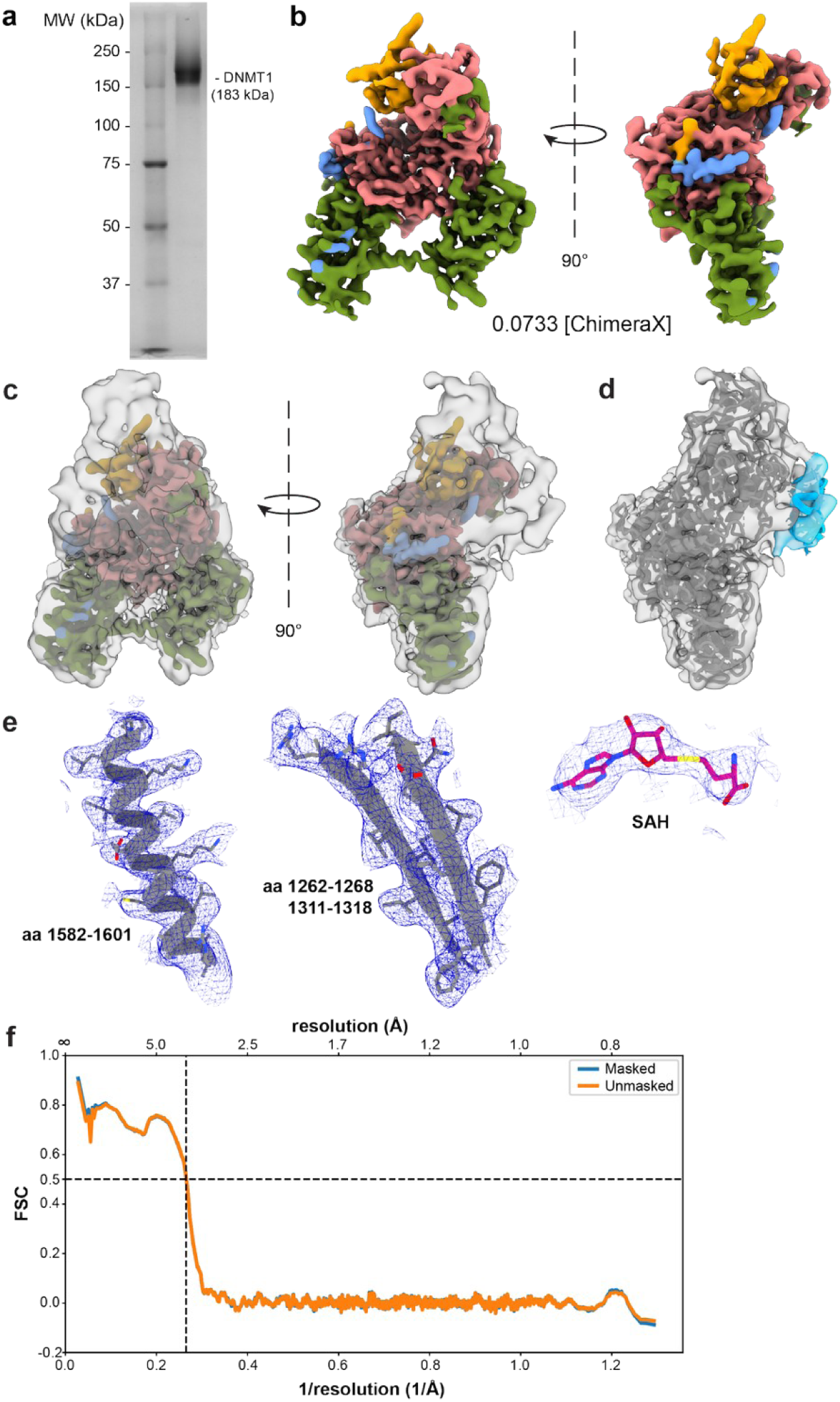
Full-length DNMT1 and pUG-fold RNA map and model details. a, Coomassie-stained SDS gel of purified full-length DNMT1 **b,** Cryo-EM density map shown at a higher volume threshold (0.0733, ChimeraX), colored by domain. **c,**Cryo-EM density map shown at two volume threshold levels. The transparent surface at a low threshold (0.0239, ChimeraX) and the solid surface colored by domain at a higher volume threshold (0.0733, ChimeraX). **d,** Additional view of the atomic models fit into the density map (PDB: 38DB). Cryo-EM density and built-in models for select regions of the final map of DNMT1 bound to GU_20_ RNA showing high-resolution details – alpha helix (left), beta sheet (middle), S-adenosyl homocysteine (SAH) (right). **e,** Fourier shell correlation (FSC) between the map and model of full-length DNMT1 with pUG-fold RNA.

**Extended Data Fig. 3:**
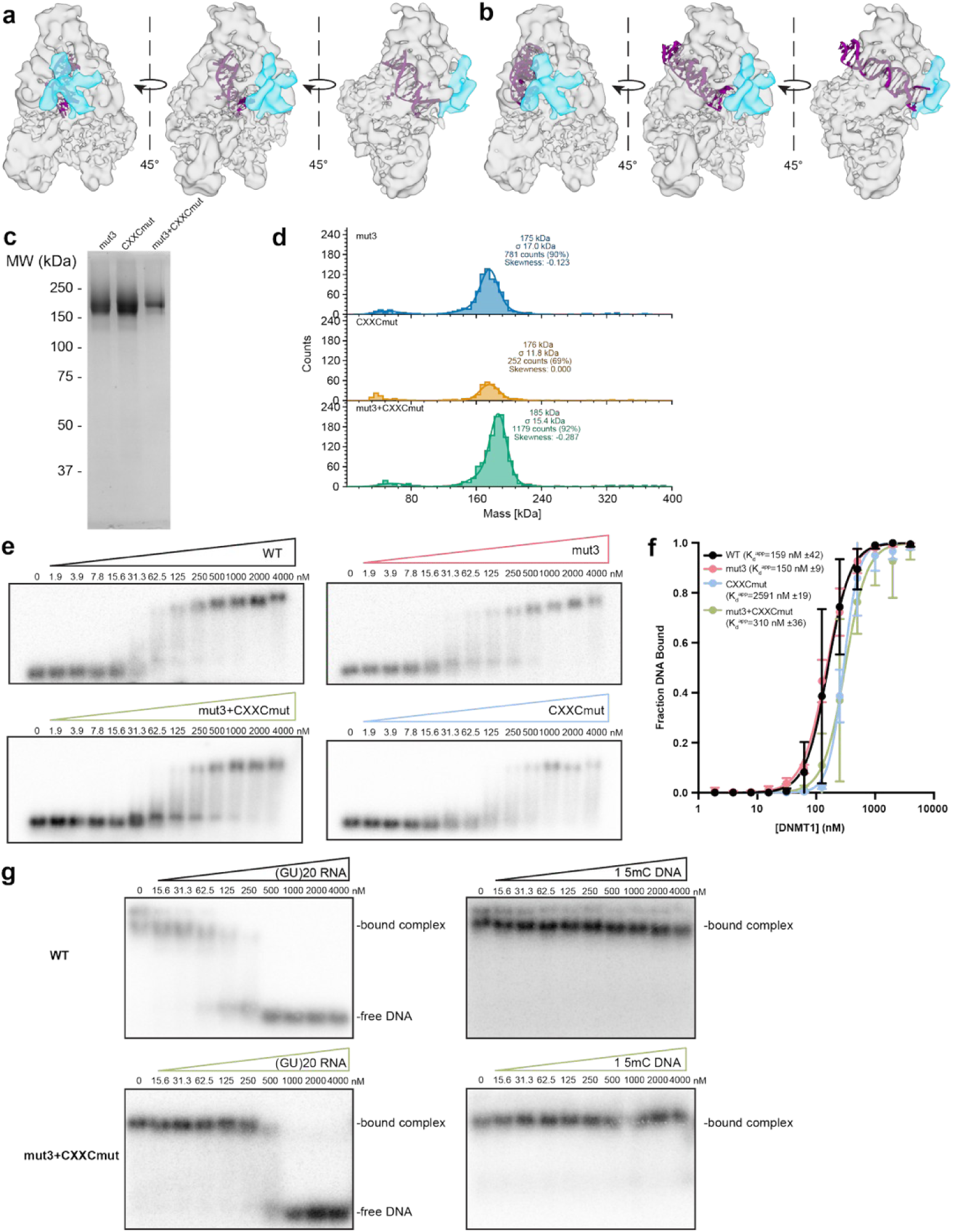
Mutations of positively charged residues of the RNA binding surface do not affect RNA binding. a, Rigid body docking of hemi-methylated DNA (purple, PDB: 4DA4) in the active site of DNMT1. **b,**Superposition of unmethylated DNA (purple, PDB: 3PTA) bound to the CXXC domain of DNMT1. **c,**Coomassie-stained SDS gel of mutant DNMT1 constructs. **d,** Mass photometry of mutant DNMT1 constructs, expected molecular weight = 183 kDa. **e,** Representative EMSAs of trace amounts of hemi-methylated DNA construct 1 5mC DNA, containing one 5mCpG site, and increasing amounts of indicated DNMT1 construct. **f,** Binding curves of DNMT1 constructs and 1 5mC DNA from E. Error bars represent standard deviation of 3 independent experiments. **g,** Representative binding competition EMSAs with trace amounts of 1 5mC DNA (left) or (GU)_20_ RNA (right) pre-incubated with either WT DNMT1 (top) or mut3+CXXCmut (bottom). Increasing amounts of (GU)_20_ RNA (left) or 1 5mC DNA (right) were added. Replicates of mut3+CXXCmut with trace amounts of 1 5mC DNA and increasing amounts in Supplementary Figure 3.

**Extended Data Fig. 4:**
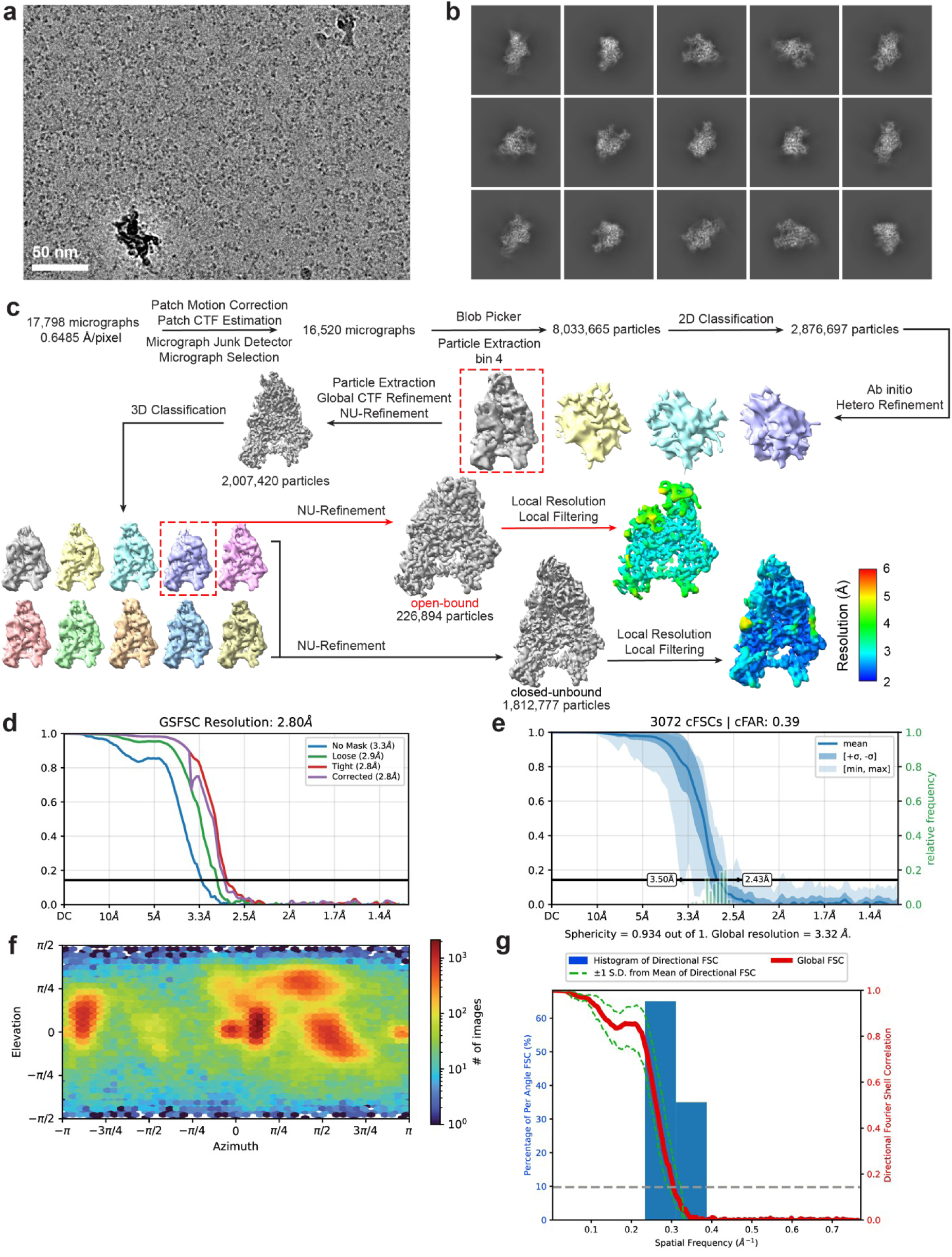
Data processing scheme for mut3+CXXCmut DNMT1 and pUG-fold RNA. a, Representative micrograph. **b,** Representative 2D projections. **c,** Data processing scheme. Red arrows show processing for the open-bound state. Black arrows show processing for the closed-bound state. Local filtered maps are colored by resolution. All data processing steps were done in Cryosparc v4.7.1. **e,** Fourier shell correlation (FSC) plot for the open-bound state. **f,** Conical Fourier shell correlation (cFSC) plot for the open-bound state. **g,** Posterior precision directional distribution plot for the open-bound state. **h,** 3D FSC plot for the open-bound state.

**Extended Data Fig. 5:**
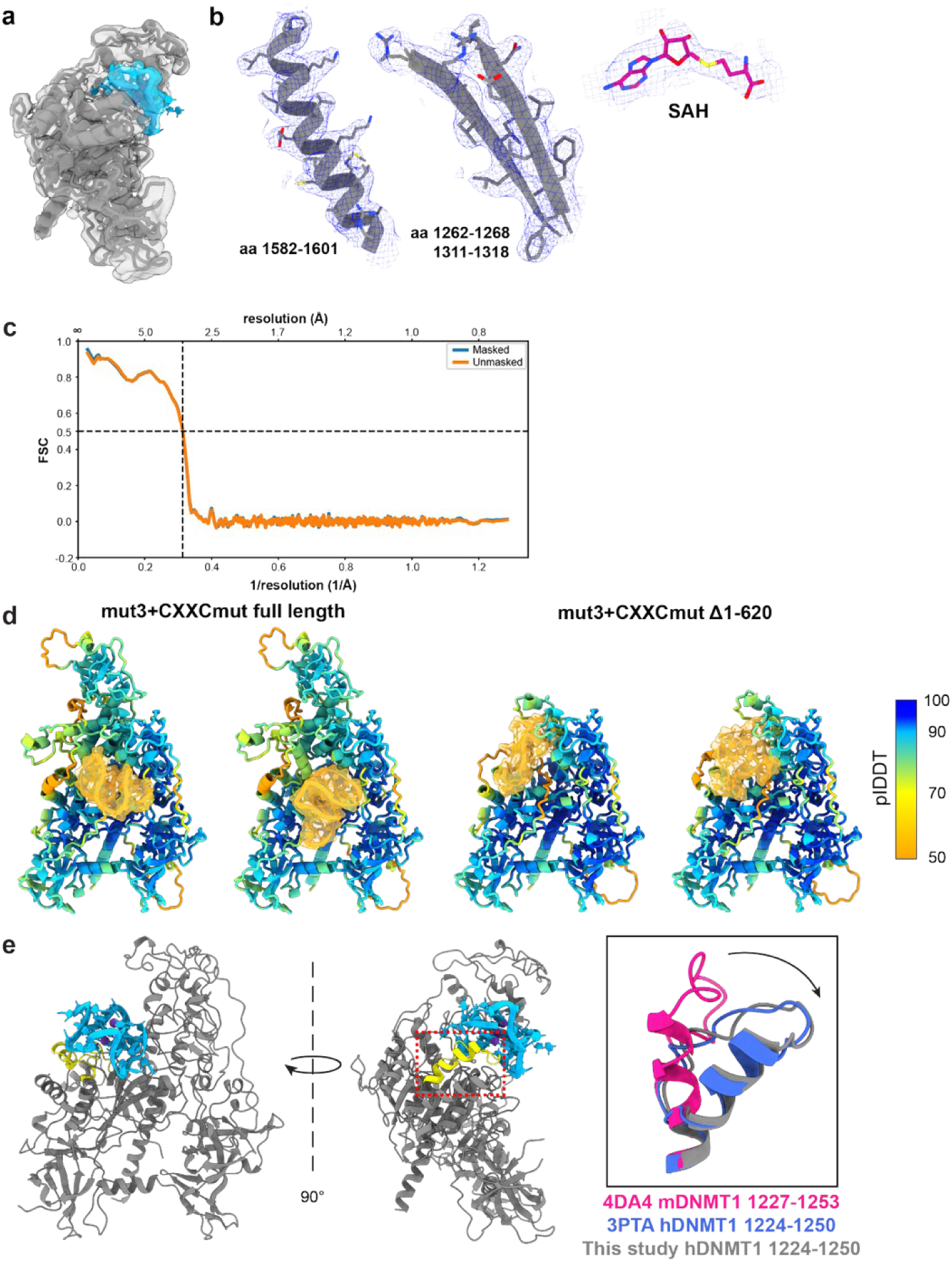
mut3+CXXCmut DNMT1 and pUG-fold RNA map and model details. a, Additional view of atomic models fit into the density map. **b,** High resolution details – alpha helix (left), beta sheet (middle), S-adenosyl homocysteine (SAH) (right). **c,** Fourier shell correlation (FSC) between the map and model of mut3+CXXCmut with pUG-fold RNA. **d,** AlphaFold3 predictions of full length mut3+CCXXCmut DNMT1 (left) with TERRA RNA ([UUAGGG]_4_, K^+^). Δ1-620 mut3+CXXCmut DNMT1 (right) with TERRA RNA ([UUAGGG]4, K^+^). Residues 1-350 omitted for clarity. Models colored by predicted Local Distance Difference Test (plDDT). Predicted aligned error (PAE) plots for models in Supplemental Figures 5-6. **e,** Position of the ⍺-helix following the catalytic loop (residues 1224-1250, yellow) in the atomic models of mut3+CXXCmut DNMT1 and pUG-fold RNA. Inset of the ⍺-helix (gray) compared to the active conformation of DNMT1 (PDB: 4DA4, pink) and inactive conformation of DNMT1 (PDB: 3PTA, royal blue), aligned by overall fits between models. Black arrow shows the movement of the catalytic loop compared to the active conformation.

**Extended Data Fig. 6:**
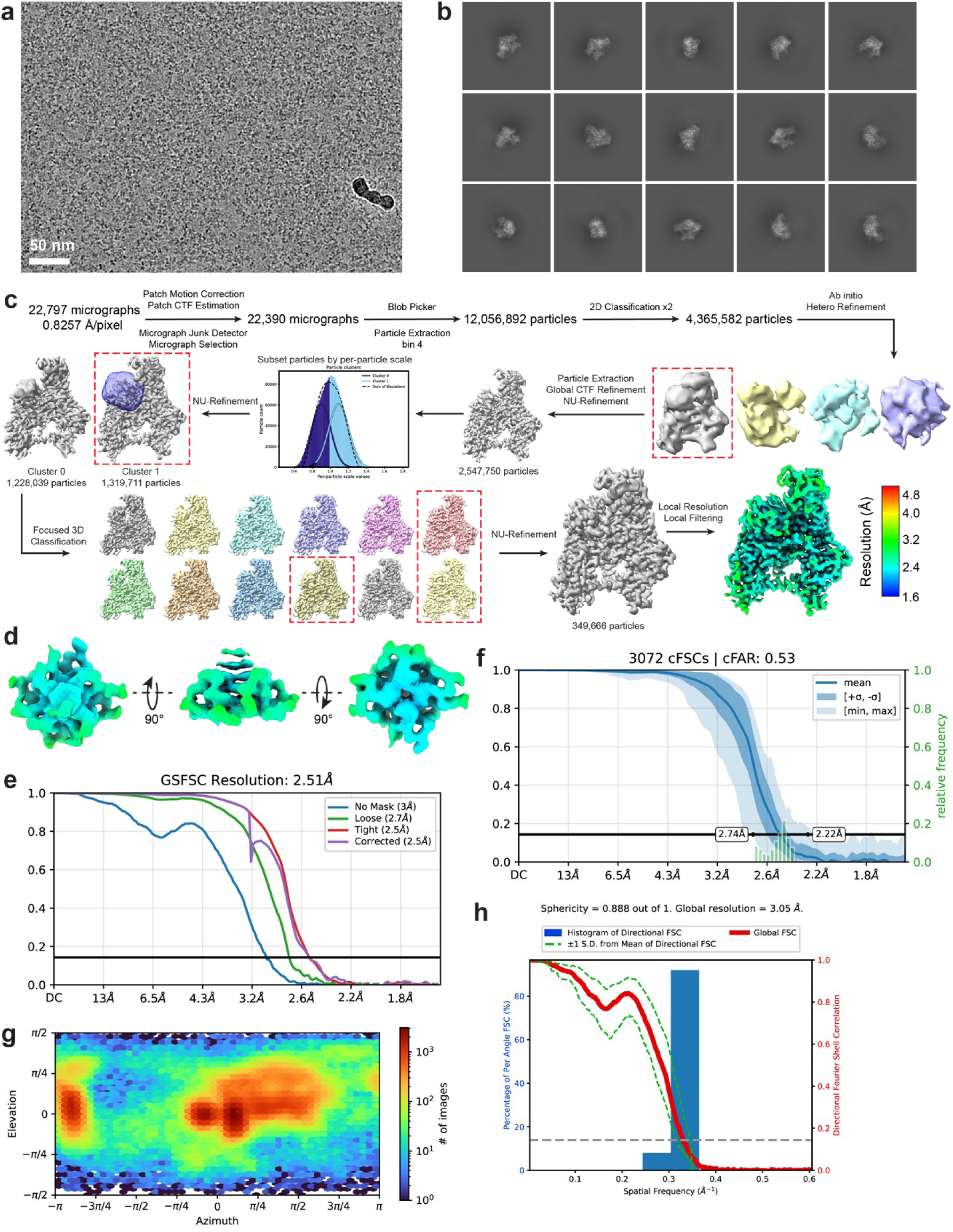
Data processing scheme for Δ1-619 DNMT1 and pUG-fold RNA. a, Representative micrograph. **b,** Representative 2D projections. **c,** Data processing scheme. Local filtered maps are colored by resolution. All data processing steps were done in Cryosparc v4.7.1. **d,** Local filtered map of the pUG-fold RNA alone, colored by resolution. **e,** Fourier shell correlation (FSC) plot. **f,** Conical Fourier shell correlation (cFSC) plot. **g,** Posterior precision directional distribution plot. **h,** 3D FSC plot.

**Extended Data Fig. 7:**
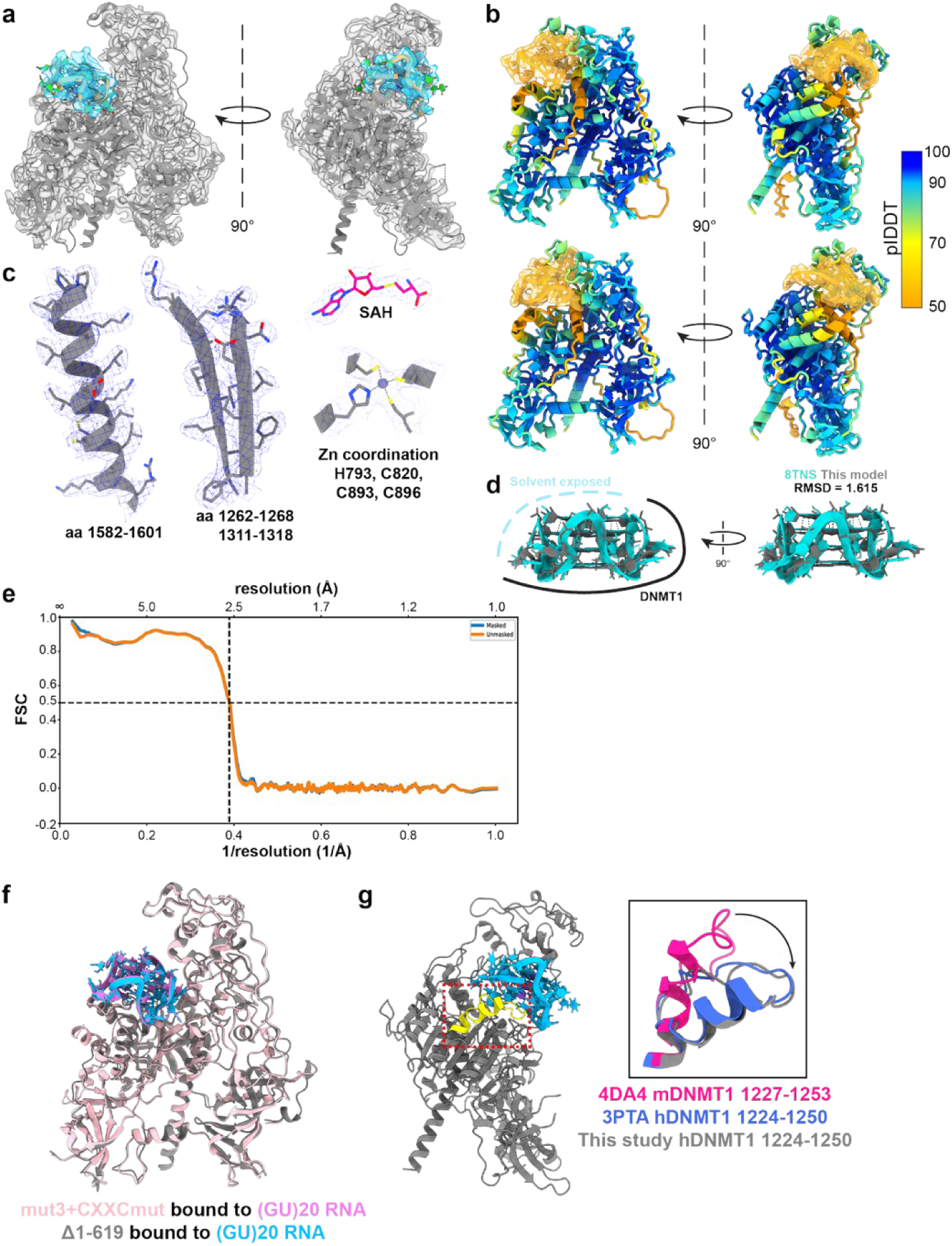
Δ1-619 DNMT1 and pUG-fold RNA map and model details. a, Atomic models fit into the density map. **b,** AlphaFold 3 predictions of Δ1-619 DNMT1 with TERRA RNA ([UUAGGG]_4_, K^+^). Models colored by predicted Local Distance Difference Test (plDDT). Predicted aligned error (PAE) plots for both models in Supplemental Figures 8-9. **c,** High-resolution details – alpha helix (left), beta sheet (middle), S-adenosyl homocysteine (SAH) (top right), and zinc ion coordination (bottom right). **d,** Superposition of atomic model of pUG RNA from this study (gray) and the NMR model (PDB: 8TNS) (aqua). Root mean square deviation (RMSD) between the models is 1.615 Å. **e,** Fourier shell correlation (FSC) between the map and model of of Δ1-619 DNMT1 with pUG-fold RNA. **f,** Superposition of mut3+CXXCmut (pink) bound to (GU)_20_ RNA (purple) and Δ1-619 DNMT1 (gray) bound to (GU)_20_ RNA (blue). **g,** Position of the ⍺-helix following the catalytic loop (residues 1224-1250, yellow) in the atomic models of Δ1-619 DNMT1 and pUG-fold RNA. Inset of the ⍺-helix (gray) compared to the active conformation of DNMT1 (PDB: 4DA4, pink) and inactive conformation of DNMT1 (PDB: 3PTA, royal blue), aligned by overall fits between models. Black arrow shows the movement of the catalytic loop compared to the active conformation.

**Extended Data Fig. 8:**
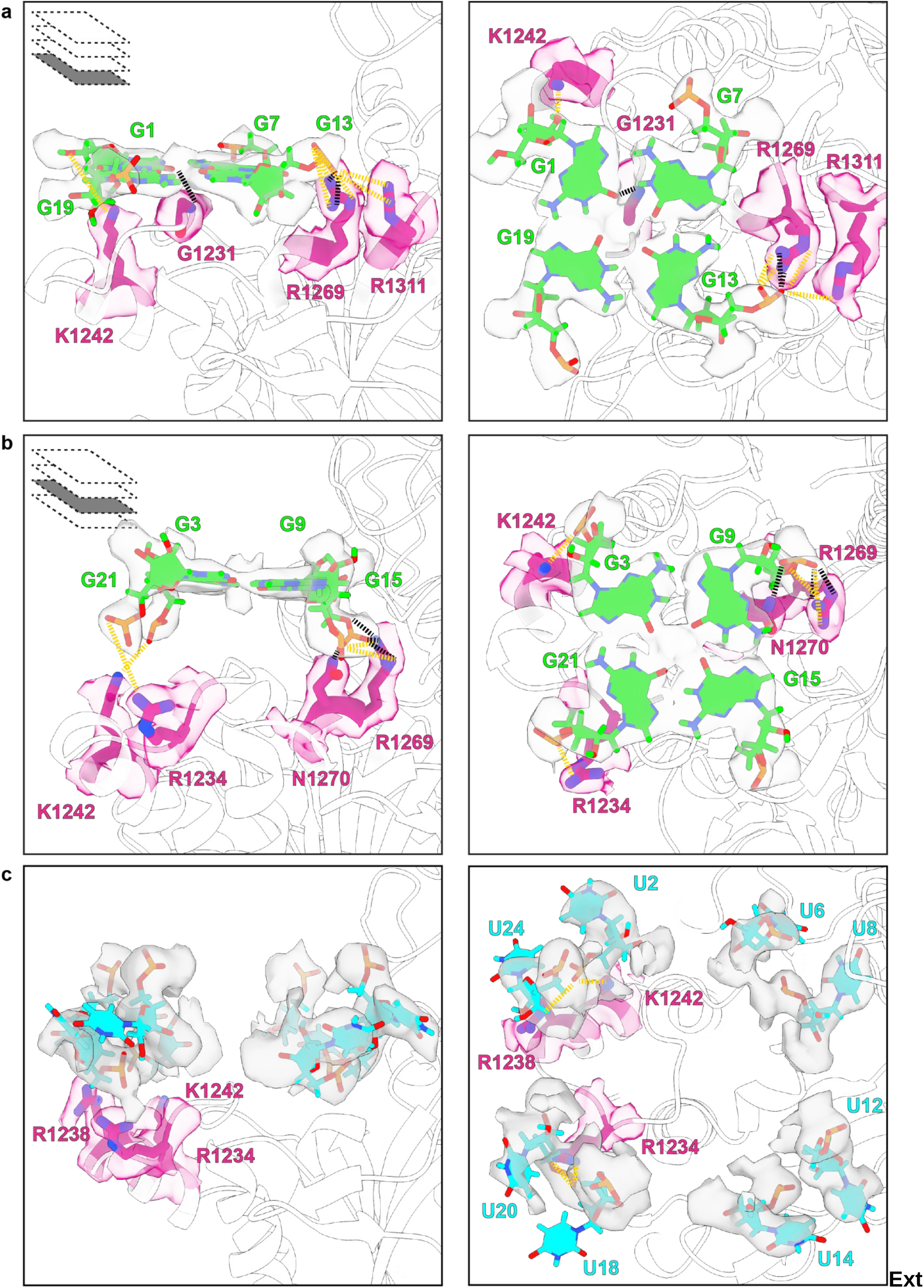
Hydrogen bond and electrostatic interactions between DNMT1 (Δ1- 619) and pUG-fold RNA. a, Views of the G1 quartet interactions with DNMT1. **b,** Views of the G3 quartet interactions with DNMT1. **c,** Views of the flipped-out uracil nucleotides’ interactions with DNMT1. Hydrogen bonds are shown in black dashed lines. Electrostatic interactions are shown in yellow dash lines. Guanine bases shown in green, uracil bases shown in cyan, phosphate groups shown in orange. Interacting protein residues shown in medium violet red.

**Extended Data Fig. 9:**
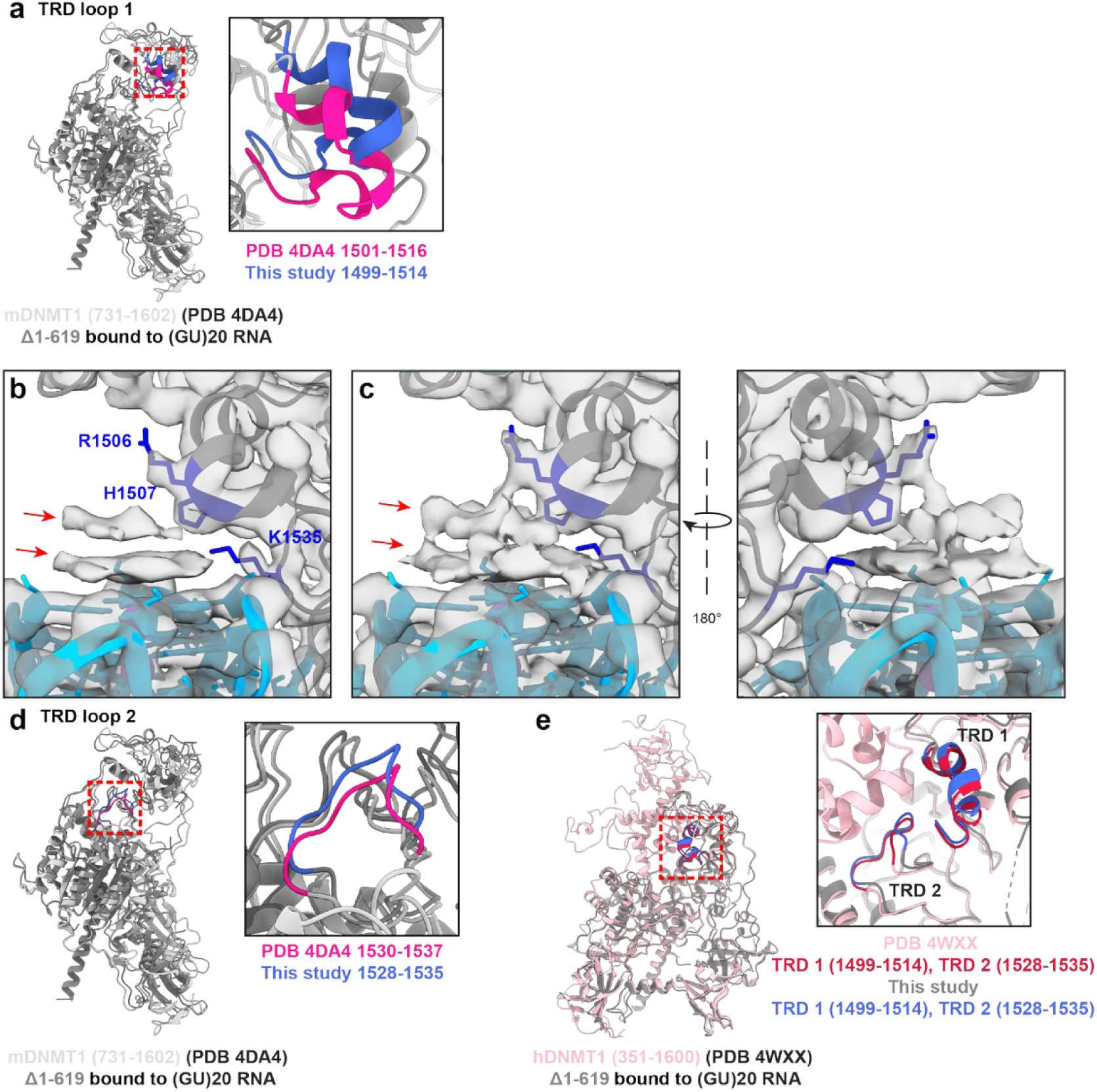
Positioning of TRD loops in DNMT1 (Δ1-619) bound to (GU)_20_ RNA. a, Position of TRD loop 1 in DNMT1 (Δ1-619) bound to (GU)_20_ RNA (gray, TRD loop 1 in royal blue) compared to active form DNMT1 (731-1602) bound to hemi-methylated DNA (silver, TRD loop 1 in deep pink), aligned by overall fits between models. Dashed red box shown zoomed in (right). **b,** Extra density in the cryo-EM map above the pUG-fold RNA, indicated with red arrows, relative to TRD loop 1 (blue). Cryo-EM map shown at volume threshold level 0.198 (ChimeraX) Cryo-EM map at a lower volume threshold (0.137, ChimeraX), showing connecting density between the extra density and the pUG-fold RNA density. **c,** Position of TRD loop 2 in DNMT1 (Δ1-619) bound to (GU)_20_ RNA (gray, TRD loop 2 in royal blue) compared to active form DNMT1 (731-1602) bound to hemi-methylated DNA (silver, TRD loop 2 in deep pink), aligned by overall fits between models. Dashed red box shown zoomed in (right). **d,** Position of TRD loops in DNMT1 (Δ1-619) bound to (GU)_20_ RNA (gray, TRD loops in royal blue) compared to apo-, autoinhibited DNMT1 (351-1600) (pink, TRD loop 1 in crimson), aligned by overall fits between models. Dashed red box shown zoomed in (right).

**Extended Data Fig. 10:**
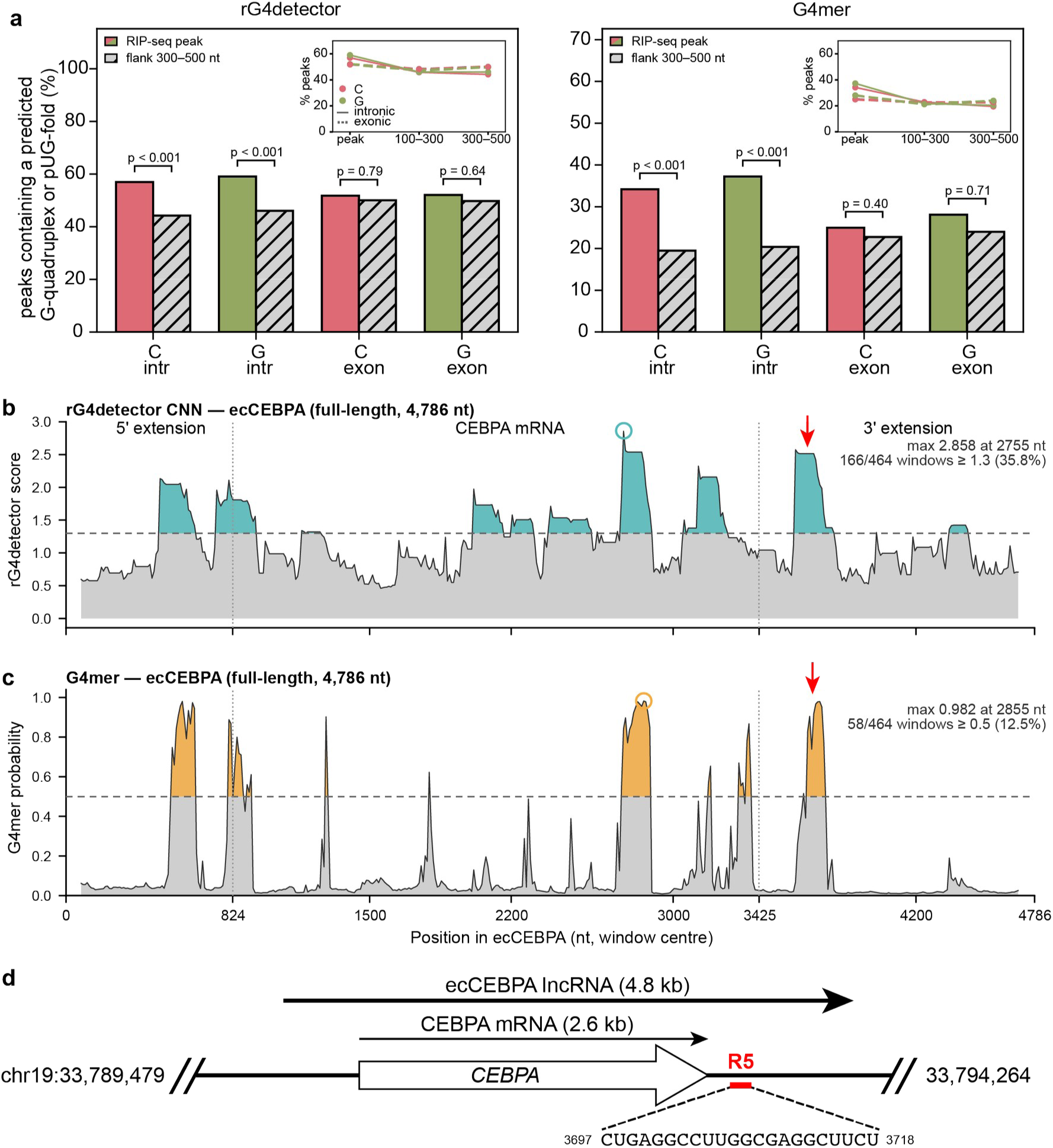
Functional differences between clusters C and G, and G- quadruplex landscapes of ecCEBPA. a, Predicted percentage of canonical or non-canonical G-quadruplexes in RIP-seq peaks versus flanking regions of the same pre-mRNA, 300–500 nt from the nearest peak center. Insets show the percentage of regions scoring positive against distance from the peak (peak, 100–300 nt, 300–500 nt) for cluster C (pink) and cluster G (green; solid lines intronic, dashed exonic). **b,** Sliding-window rG4detector score along the full length of ecCEBPA (150-nt windows, 10-nt step). **c,** Sliding-window G4mer probability score along the full length of ecCEBPA (150-nt windows, 10-nt step). In **b** and **c**, dashed horizontal line marks each method’s decision threshold. Dashed vertical lines separate the 5’ extension, CEBPA mRNA, and 3’ extension. Colored circle marks the highest scored step of the lncRNA. Red arrow represents location of R5. **d,** Diagram of CEBPA transcripts and position and sequence of R5 RNA oligonucleotide, previously shown to bind to DNMT1.

**Supplemental Figure 1.**
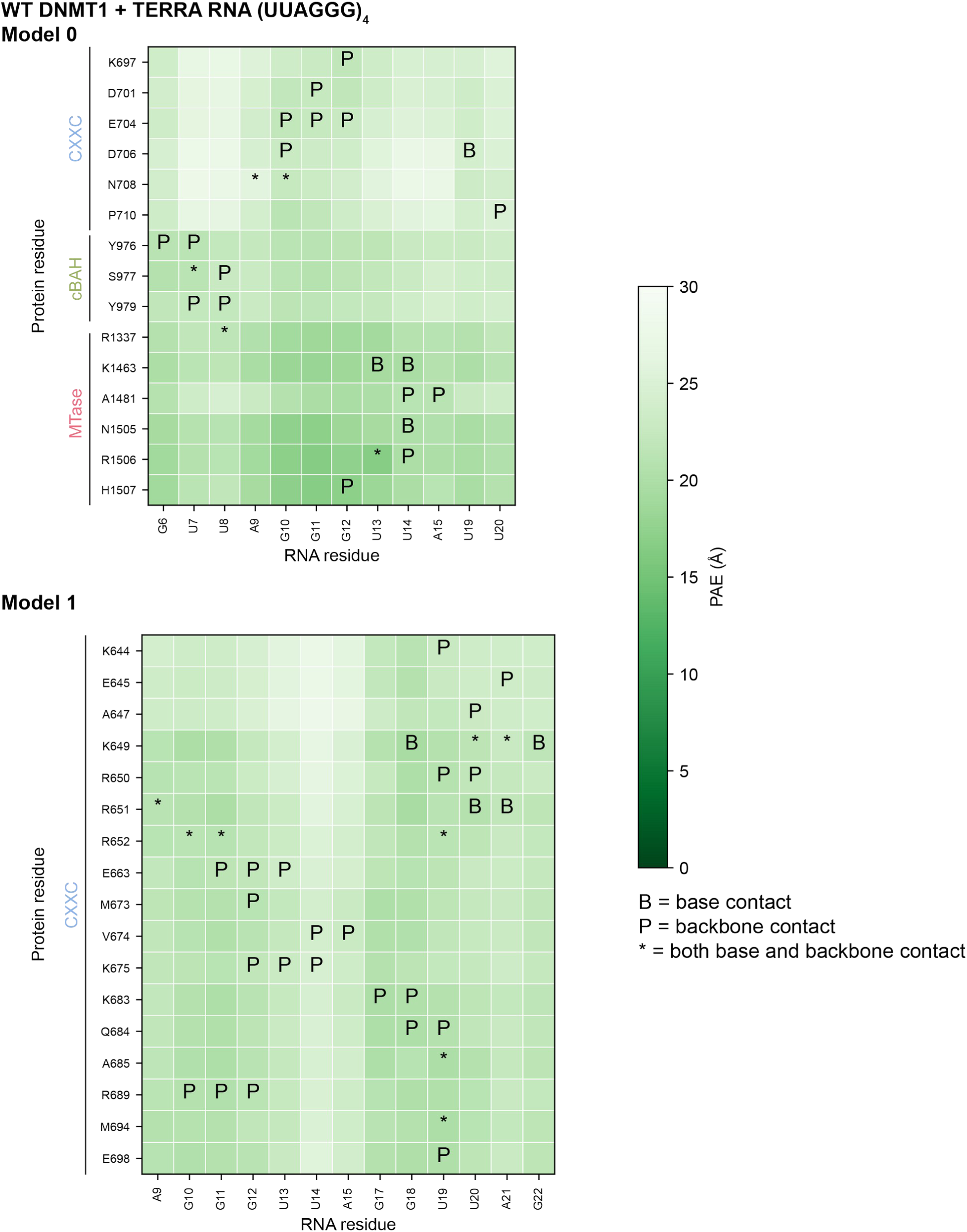
Predicted aligned error (PAE) plots for wild-type full-length DNMT1 and TERRA RNA ([UUAGGG]_4_, K^+^). Predicted align error (PAE) heat map plots for wild-type full-length DNMT1 and TERRA RNA ([UUAGGG]_4_, K^+^). Predicted interactions between protein and RNA residues. B = contact with RNA base, P = contact with phosphate backbone, * = contact with both base and phosphate backbone. Predicted models in Figure 2C.

**Supplemental Figure 2.**
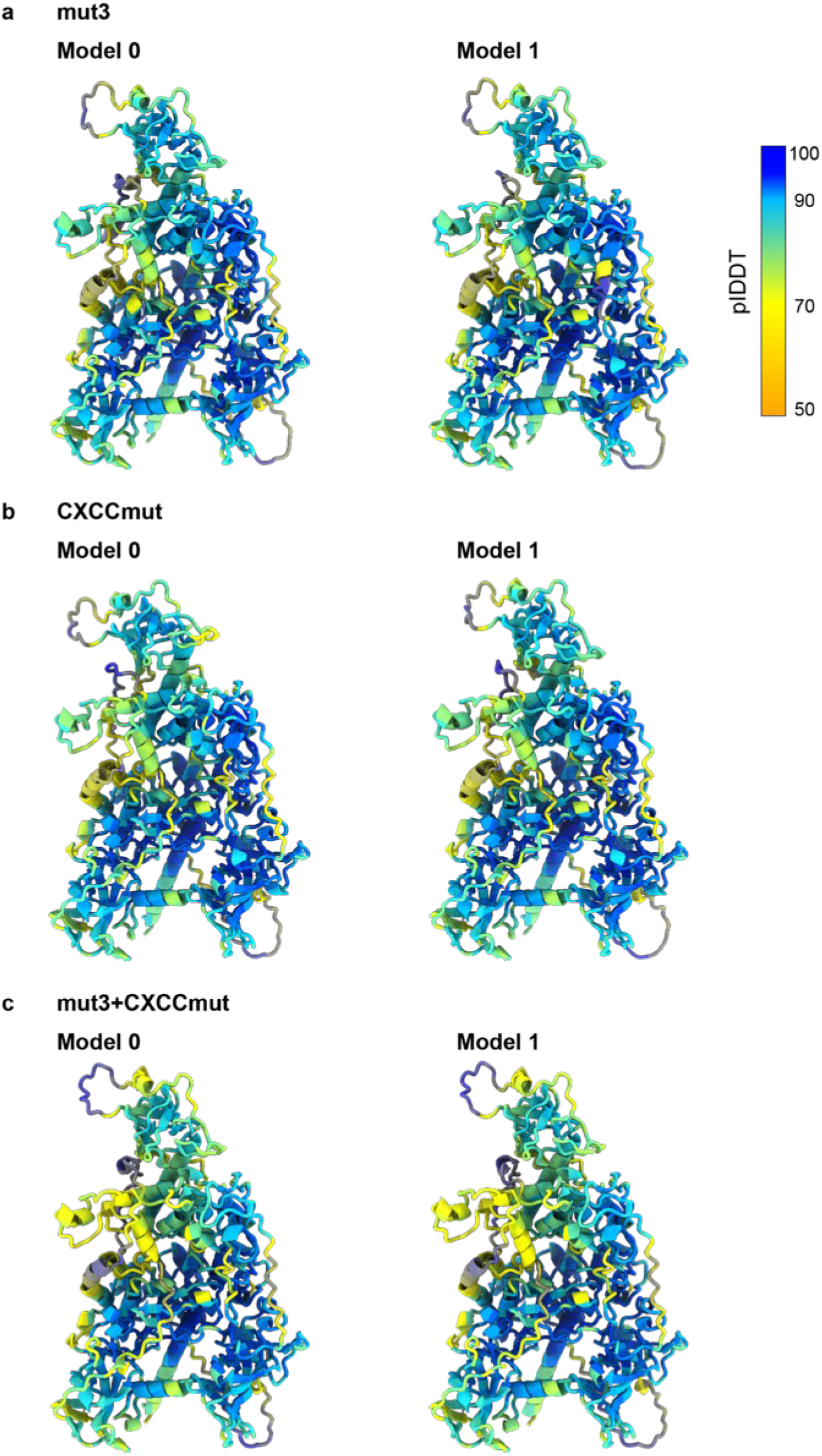
AlphaFold3 predictions of DNMT1 mutants. Predicted models for (a) mut3 (b) CXXCmut (c) mut3+CXXCmut. All models colored by predicted Local Distance Difference Test (plDDT).

**Supplemental Figure 3.**
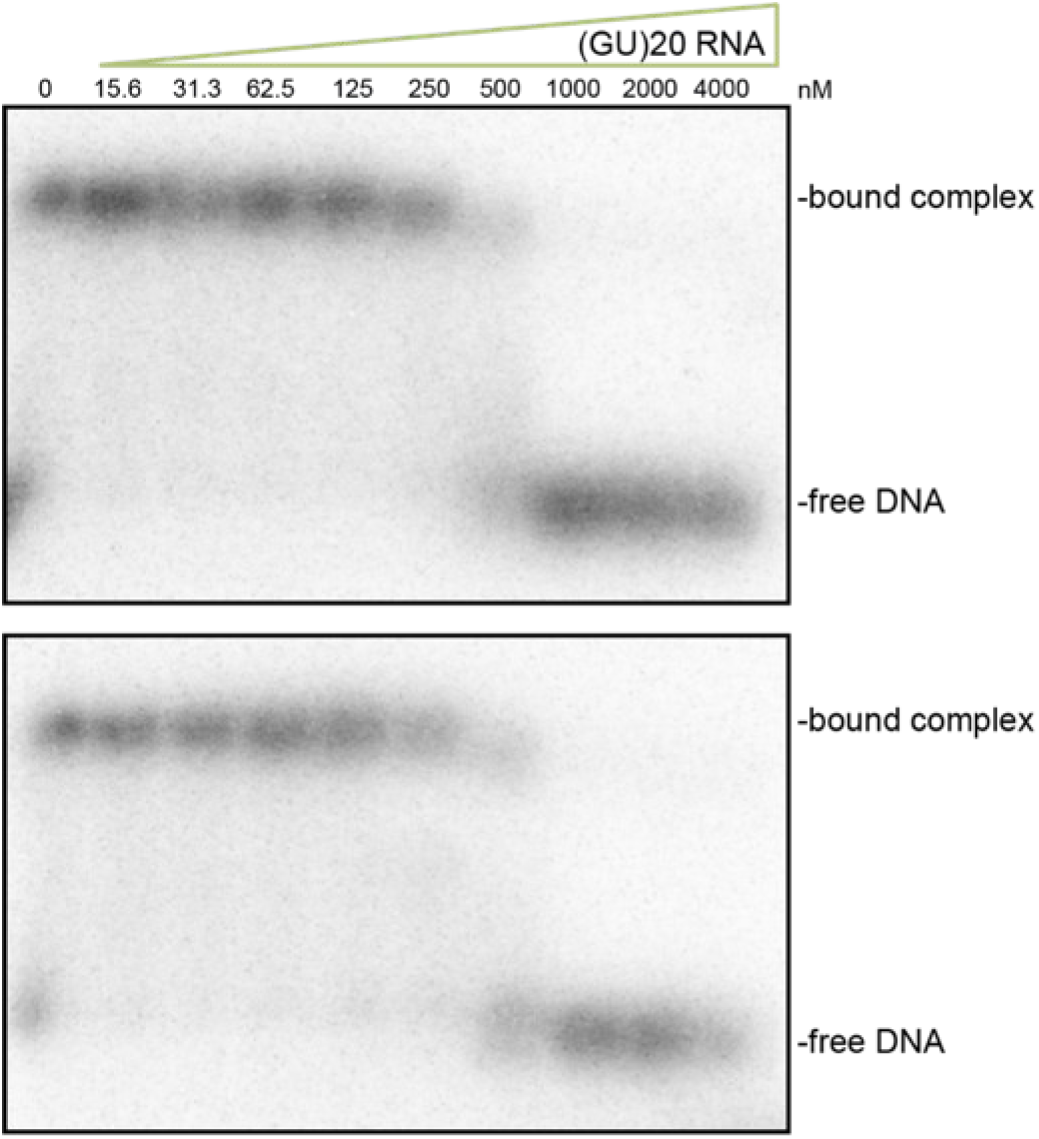
Binding competition EMSA replicates. Two independent replicates of binding competition EMSAs with trace amounts of 1 5mC DNA pre-incubated with mut3+CXXCmut. Increasing amounts of (GU)_20_ RNA were added.

**Supplemental Figure 4.** Rigid body docking of DNMT1 models into mut3+CXXCmut cryo-EM maps. (a) Open, inactive conformation of DNMT1 (646-1600) bound to non-methylated DNA (DNA not shown, PDB: 3PTA) rigid body docked into the open-bound map of mut3+CXXCmut. (b) Open, inactive conformation of DNMT1 (646-1600) bound to Zebularine- containing 12mer dsDNA (DNA not shown, PDB: 6X9I) rigid body docked into the open-bound map of mut3+CXXCmut. (c) Open, active conformation of DNMT1 (731-1600) bound to hemi-methylated DNA (DNA not shown, PDB: 4DA4) rigid body docked into the open-bound map of mut3+CXXCmut. (d) Apo-, autoinhibited conformation of DNMT1 (351-1600, PDB: 4WXX) rigid body docked into the open-bound map of mut3+CXXCmut to demonstrate the absence of most of the RFTS domain and the flexible region of the CXXC domain. (e) Apo-, autoinhibited conformation of DNMT1 (351-1600, PDB: 4WXX) rigid body docked into the closed-unbound map of mut3+CXXCmut. (f) Full-length DNMT1 bound to pUG-fold RNA (RNA not shown, from this study) rigid body docked into the closed-unbound map of mut3+CXXCmut.

**Supplemental Figure 5.**
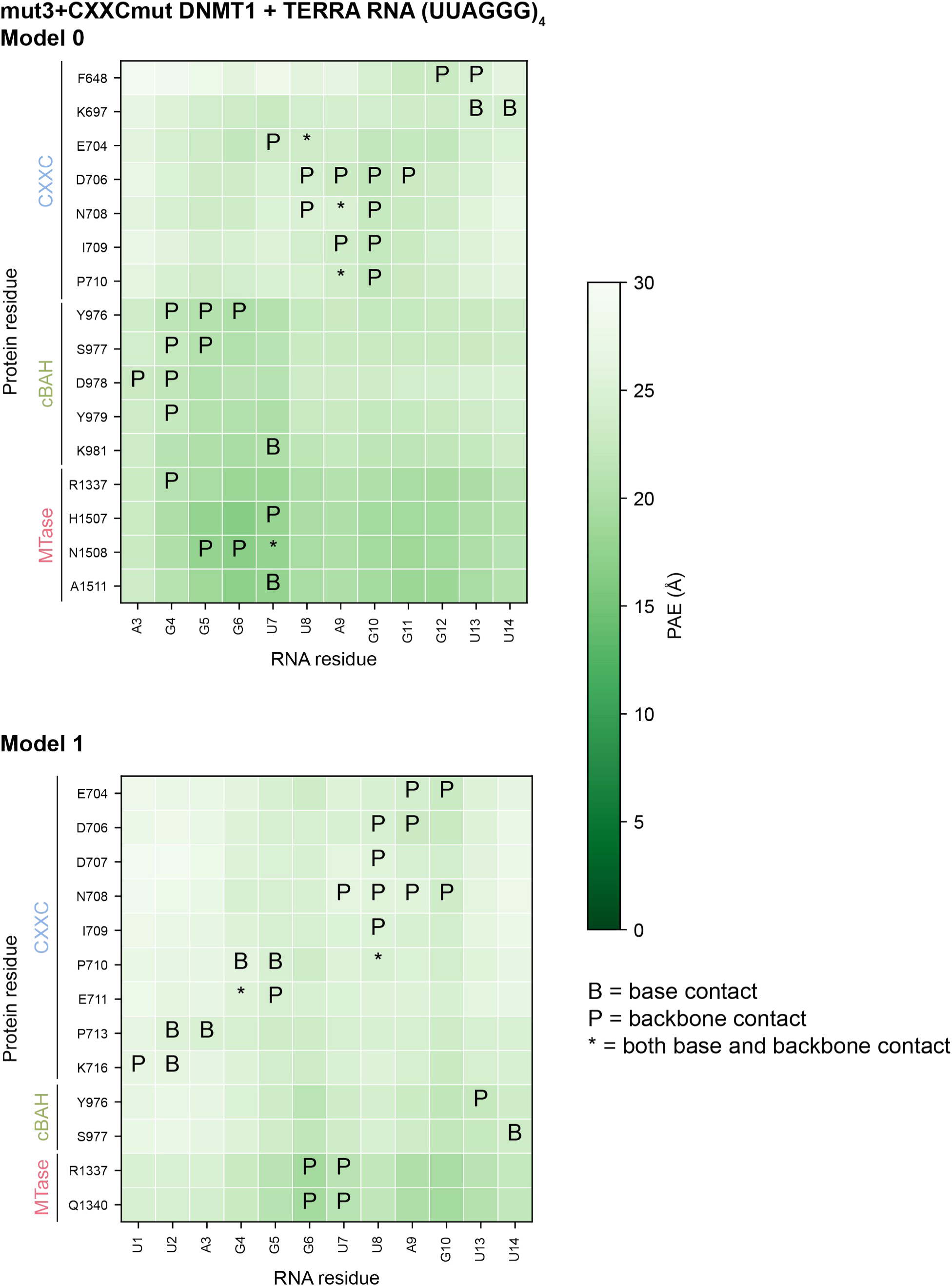
Predicted aligned error (PAE) plots for mut3+CXXCmut DNMT1 and TERRA RNA ([UUAGGG]_4_, K^+^). Predicted align error (PAE) heat map plots for mut3+CXXCmut DNMT1 and TERRA RNA ([UUAGGG]_4_, K^+^). Predicted interactions between protein and RNA residues. B = contact with RNA base, P = contact with phosphate backbone, * = contact with both base and phosphate backbone. Predicted models shown in Extended Data Figure 5D.

**Supplemental Figure 6.**
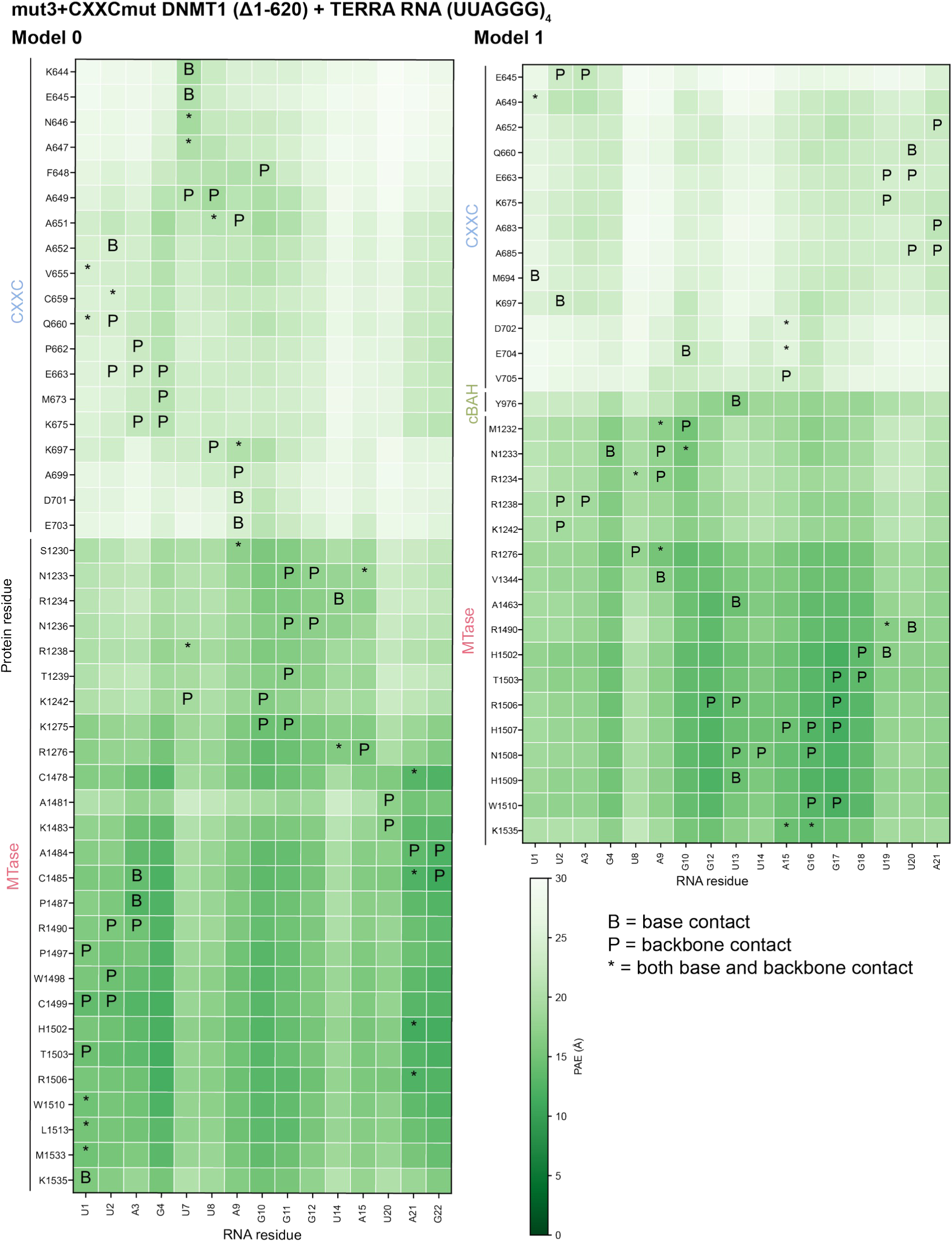
Predicted aligned error (PAE) plots for mut3+CXXCmut DNMT1 (Δ1-620) and TERRA RNA ([UUAGGG]_4_, K^+^). Predicted align error (PAE) heat map plots for mut3+CXXCmut DNMT1 (Δ1-620) and TERRA RNA ([UUAGGG]_4_, K^+^). Predicted interactions between protein and RNA residues. B = contact with RNA base, P = contact with phosphate backbone, * = contact with both base and phosphate backbone. Predicted models shown in Extended Data Figure 5D.

**Supplemental Figure 7.**
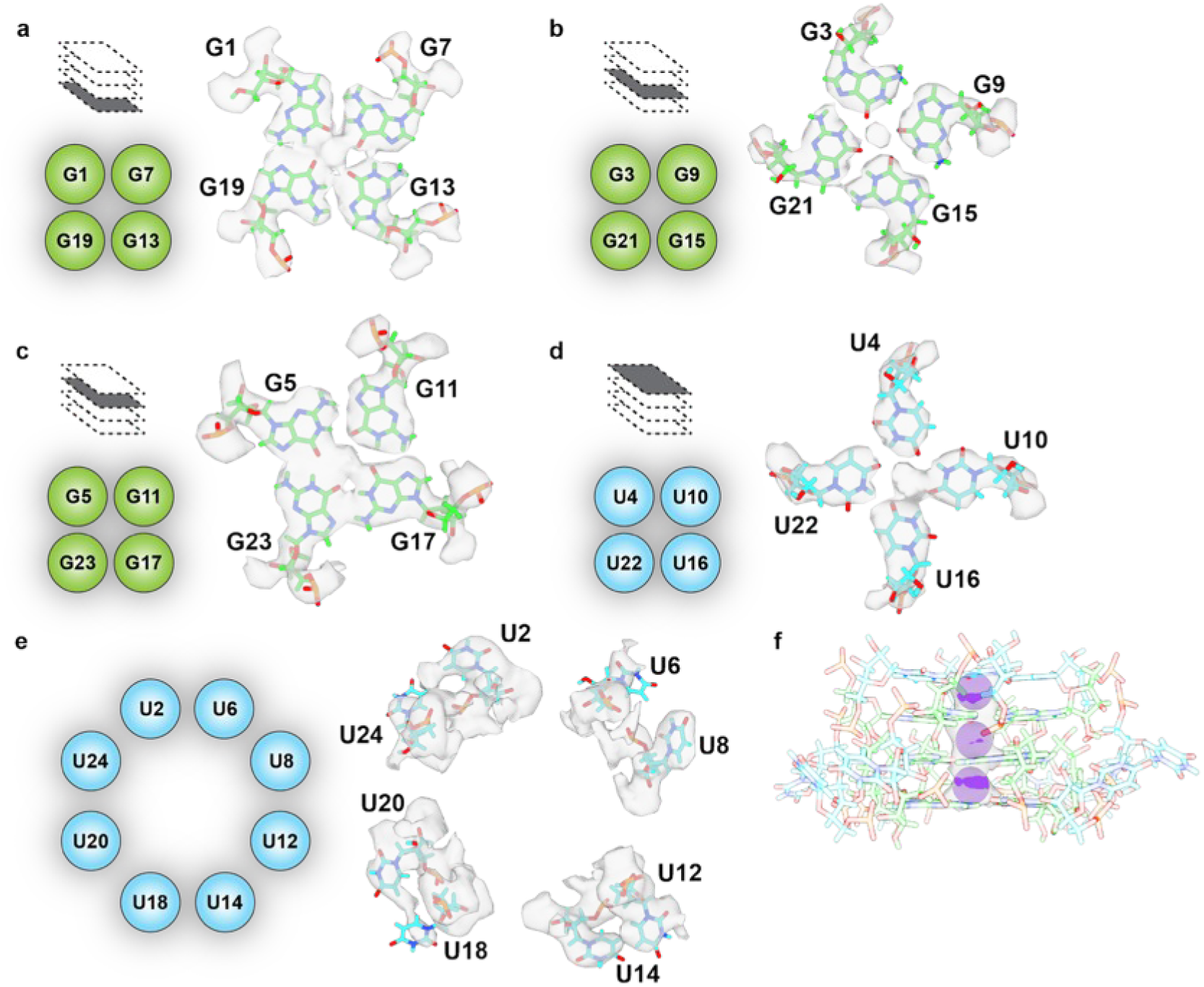
pUG-fold RNA details. (a) G1 tetrad schematic (left) and the atomic model fit into the cryo-EM density (right). (b) G3 tetrad schematic (left) and the atomic model fit into the cryo-EM density (right). (c) G5 tetrad schematic (left) and the atomic model fit into the cryo-EM density (right). (d) U tetrad schematic (left) and the atomic model fit into the cryo-EM density (right). (e) Flipped-out uracil nucleotides (left) and the atomic model fit into the cryo-EM density (right). (f) Potassium ions (K^+^) fit (purple) fit into the cryo-EM density. In all panels, guanine nucleotides shown in lime, uracil nucleotides in cyan, and phosphates in orange.

**Supplemental Figure 8.**
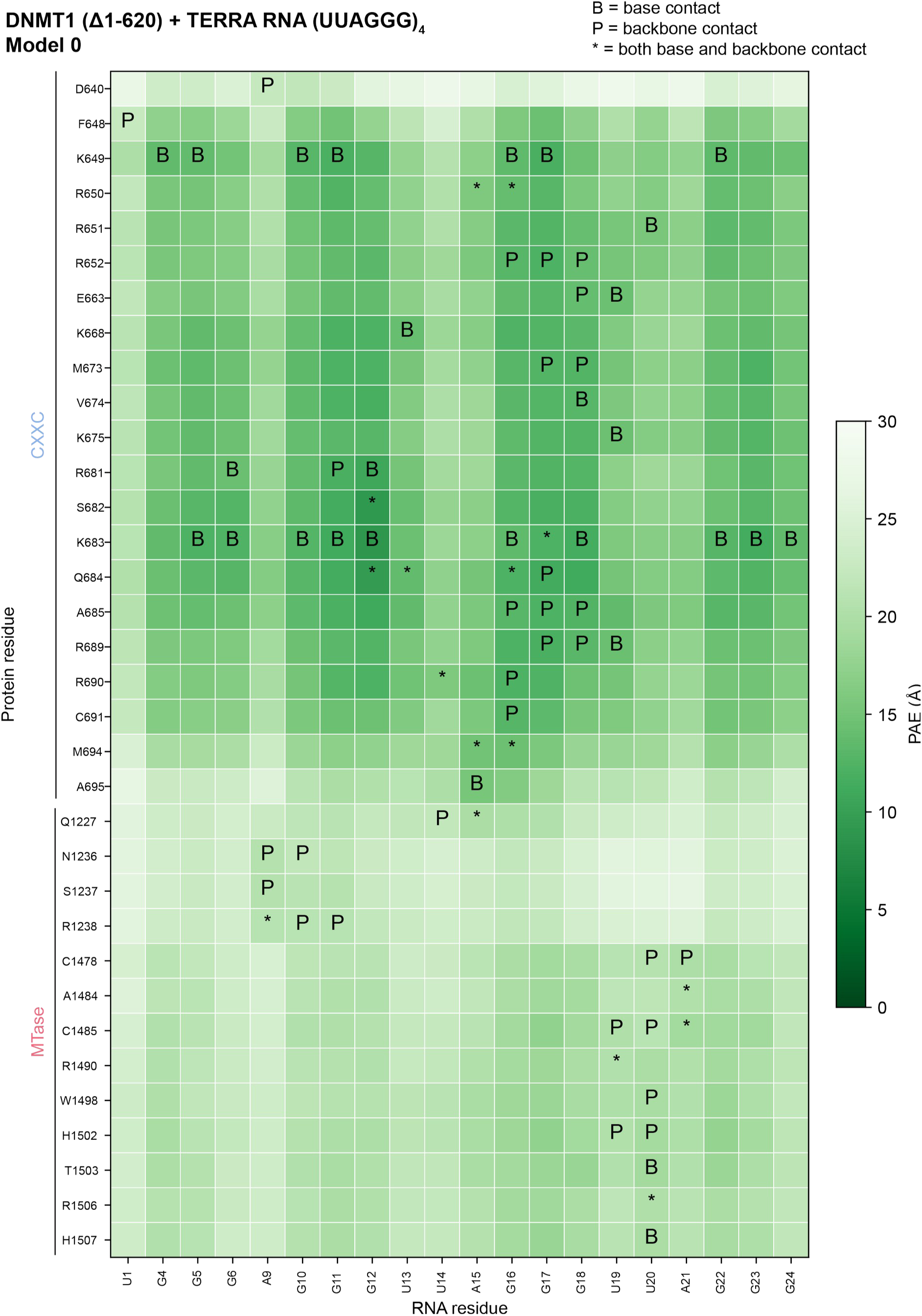
Predicted aligned error (PAE) plot for DNMT1 (Δ1-620) and TERRA RNA ([UUAGGG]_4_, K^+^) Model 0. Predicted align error (PAE) heat map plot for DNMT1 (Δ1-620) and TERRA RNA ([UUAGGG]_4_, K^+^) Model 0. Predicted interactions between protein and RNA residues. B = contact with RNA base, P = contact with phosphate backbone, * = contact with both base and phosphate backbone. Predicted models shown in Extended Data Figure 7B.

**Supplemental Figure 9.**
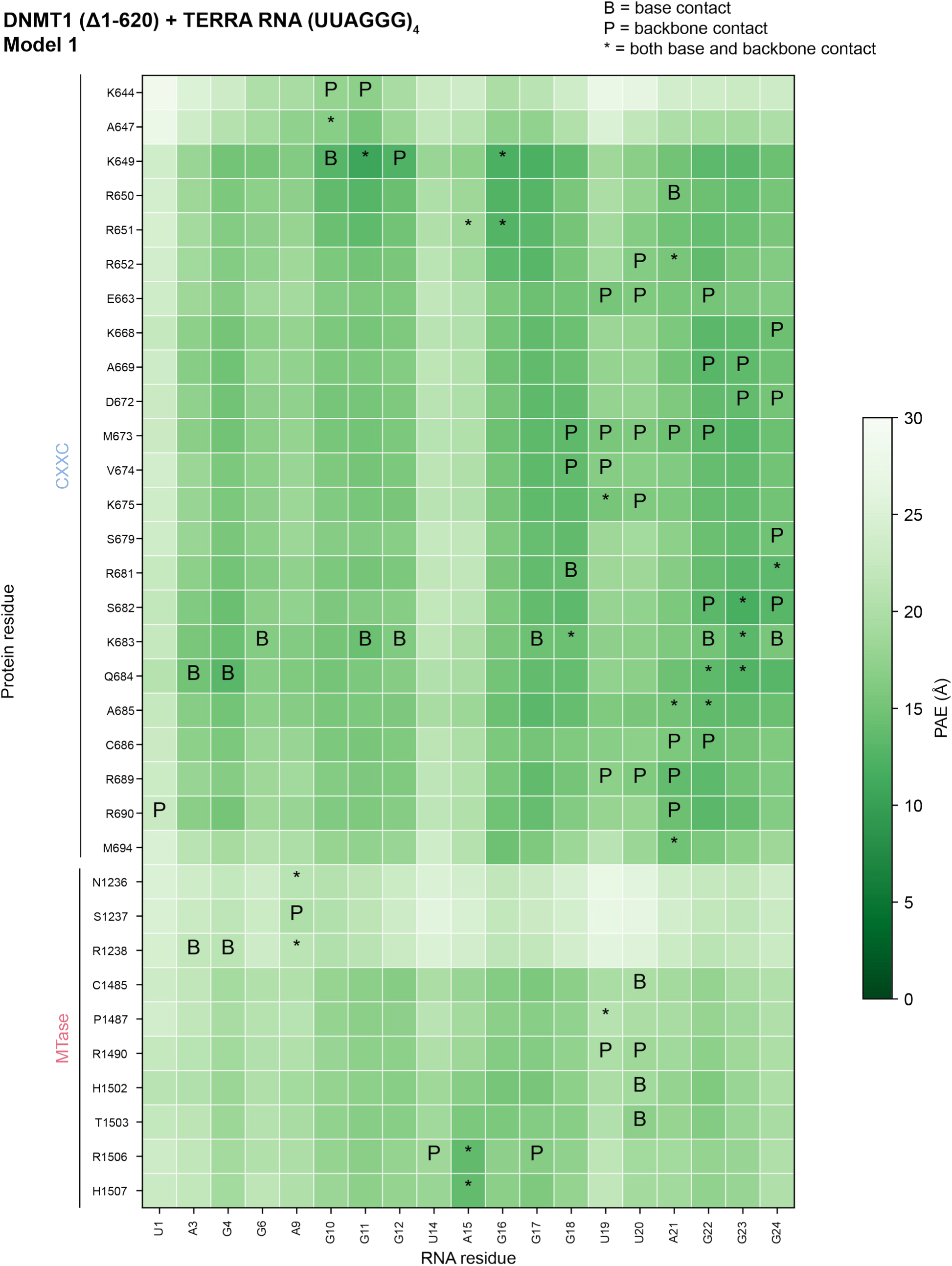
Predicted aligned error (PAE) plot for DNMT1 (Δ1-620) and TERRA RNA ([UUAGGG]_4_, K^+^) Model 1. Predicted align error (PAE) heat map plot for DNMT1 (Δ1-620) and TERRA RNA ([UUAGGG]_4_, K^+^) Model 1. Predicted interactions between protein and RNA residues. B = contact with RNA base, P = contact with phosphate backbone, * = contact with both base and phosphate backbone. Predicted models shown in Extended Data Figure 7B.

**Supplemental Figure 10.**
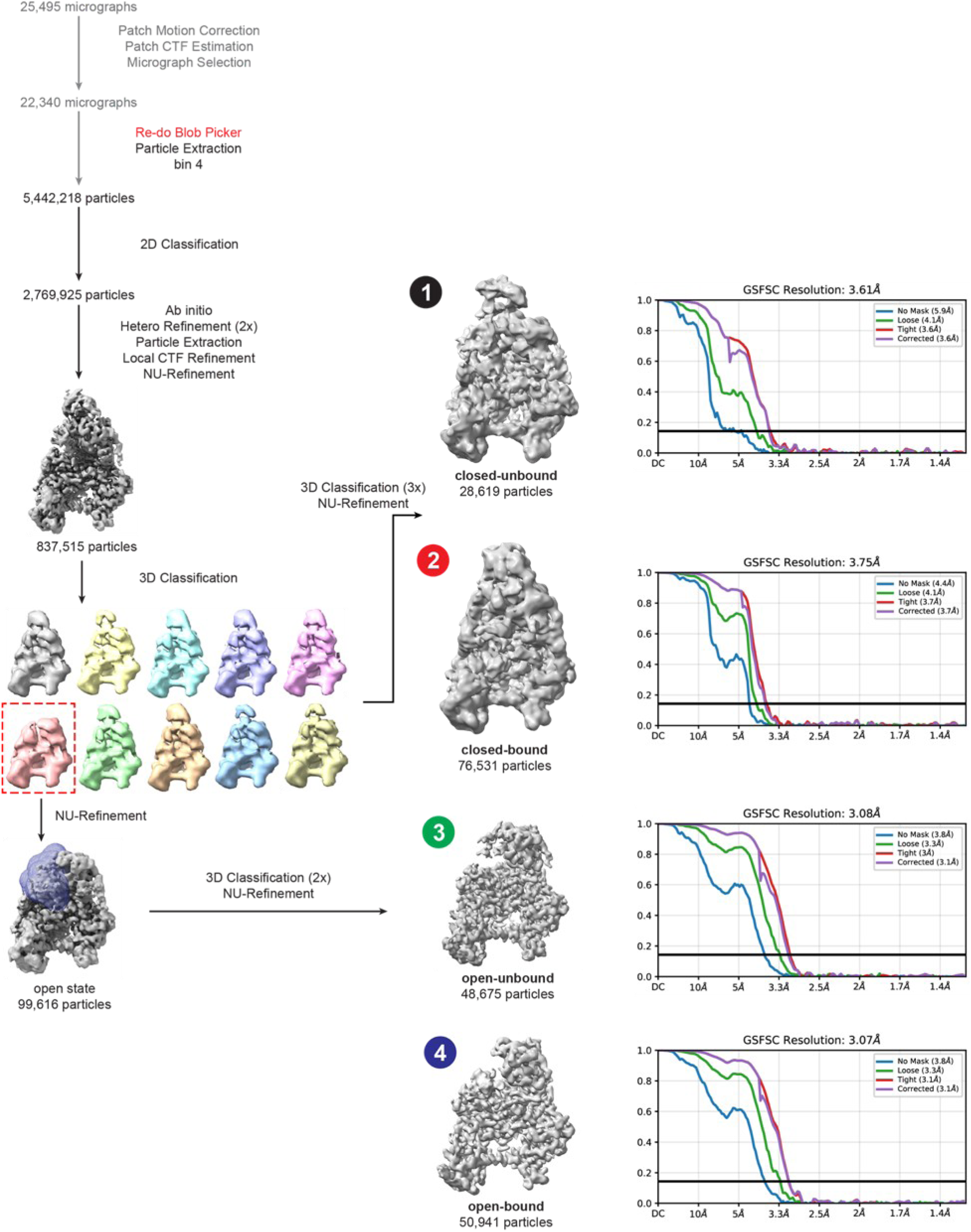
Reanalysis of data processing pipeline for wild-type full-length DNMT1 and (GU)_20_ RNA. Reanalysis of the wild-type full-length DNMT1 and GU)20 RNA yielded four conformation states: (1) closed-unbound, (2) closed-bound, (3) open-unbound, and (4) open-bound.

**Supplemental Figure 11.**
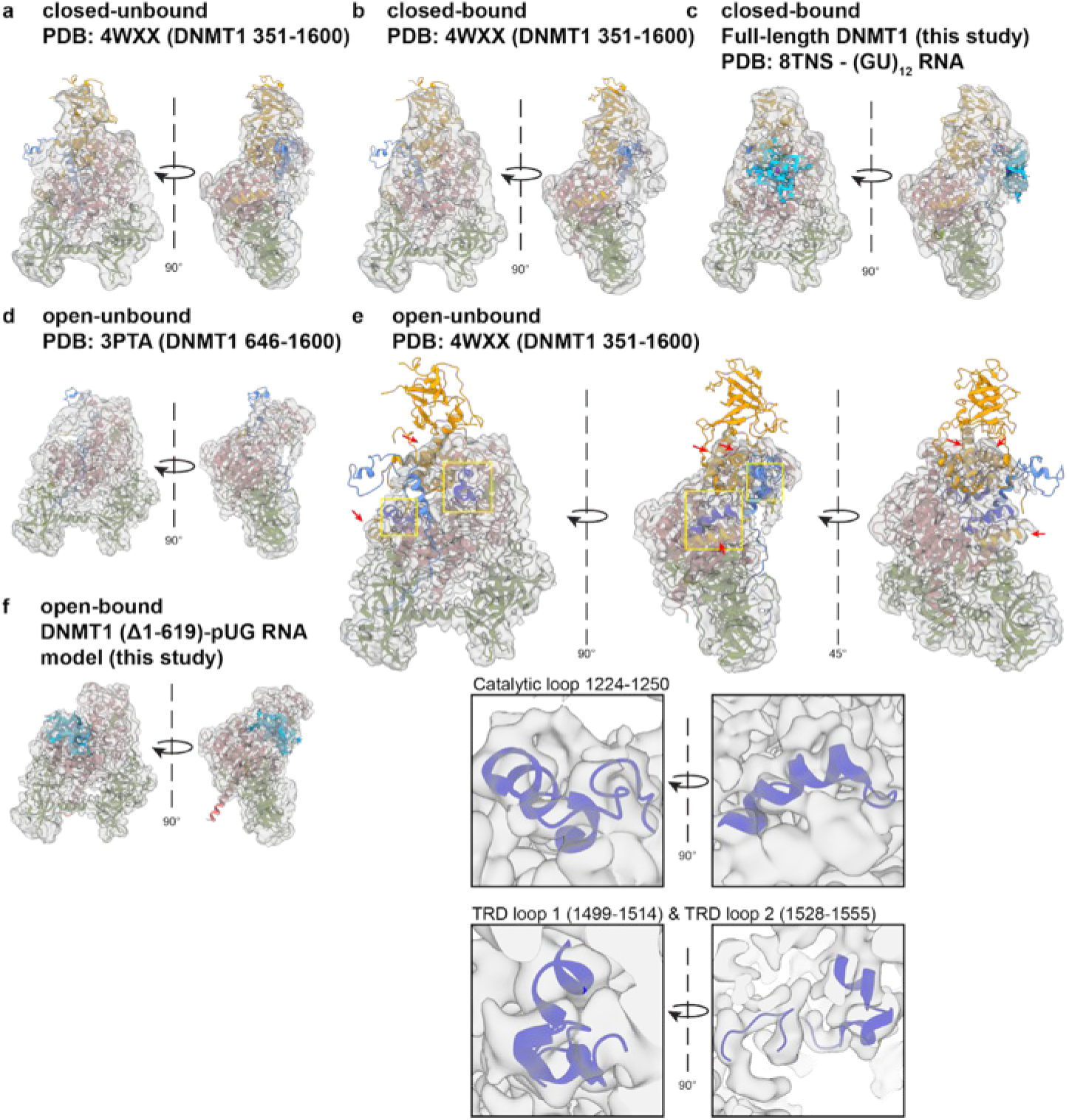
Rigid body docking of DNMT1 models into wild-type full length DNMT1 and (GU)_20_ RNA maps. (a) Apo-, autoinhibited conformation of DNMT1 (351-1600, PDB: 4WXX) rigid body docked into the closed-unbound map. (b) Apo-, autoinhibited conformation of DNMT1 (351-1600, PDB: 4WXX) rigid body docked into the closed-bound map. (c) Full-length DNMT1 bound to pUG-fold RNA (RNA not shown, from this study) and pUG-fold RNA (PDB: 8TNS) rigid body docked into the closed-bound map. (d) Open, inactive conformation of DNMT1 (646-1600) bound to non-methylated DNA (DNA not shown, PDB: 3PTA) rigid body docked into the open-unbound map. (e) Apo-, autoinhibited conformation of DNMT1 (351-1600, PDB: 4WXX) rigid body docked into the open-unbound map. Red arrows show the RFTS helices present in this conformation. Yellow boxes highlight the catalytic loop and TRD loops and the view of insets below. (f) DNMT1 (Δ1-619) bound to pUG-fold RNA (this study) rigid body docked into the open-bound map.

**Supplemental Figure 12.**
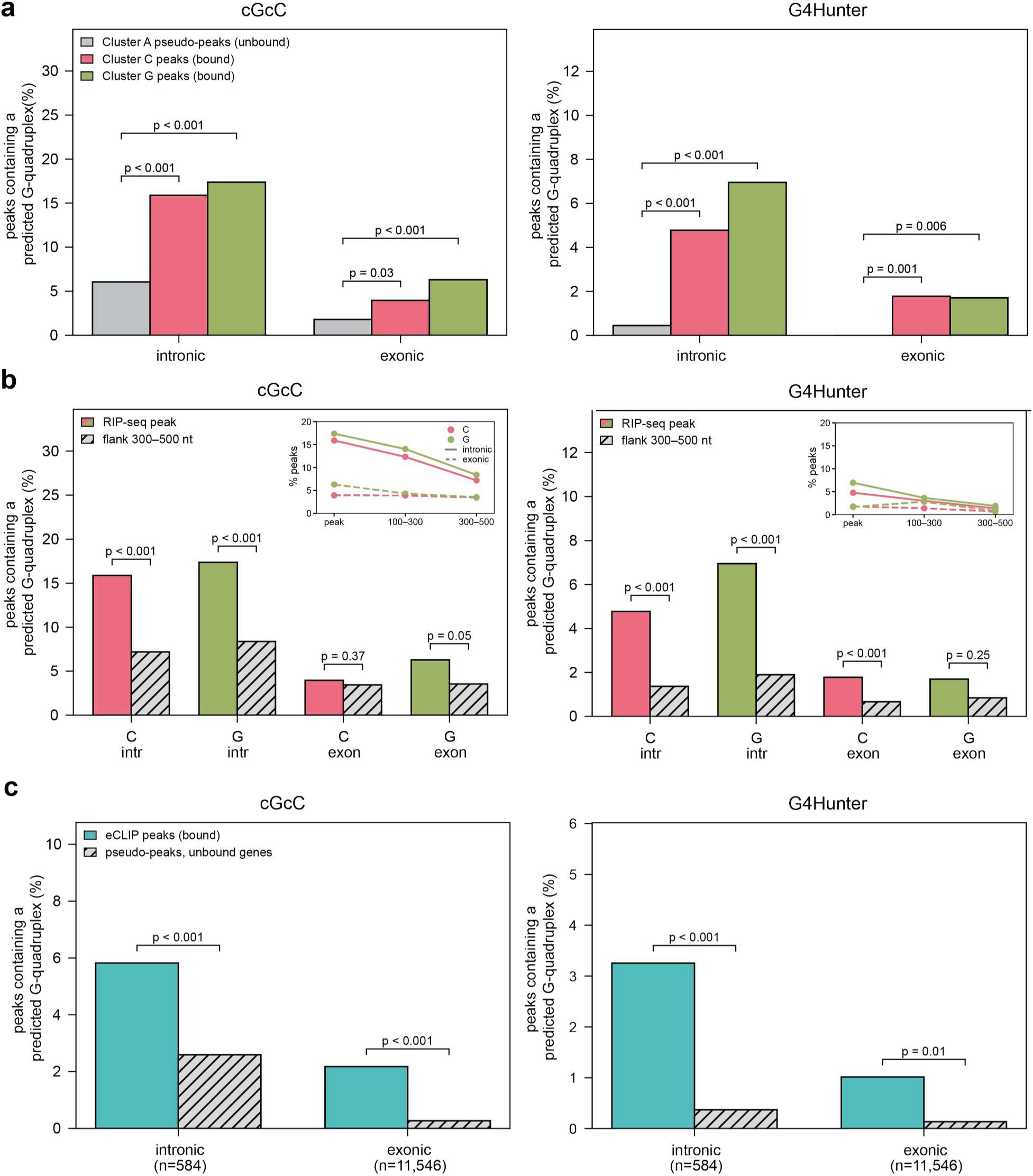
Rule-based prediction of classical G-quadruplexes in RIP-seq and eCLIP data. (a) Predicted percentage of canonical or non-canonical G-quadruplexes in RIP-seq peaks in cluster C and cluster G, versus pseudo-peaks placed at random positions in the pre-mRNA of cluster A genes. (b)Predicted percentage of canonical or non-canonical G-quadruplexes in RIP-seq peaks versus flanking regions of the same pre-mRNA, 300–500 nt from the nearest peak center. Insets show the percentage of regions scoring positive against distance from the peak (peak, 100–300 nt, 300–500 nt) for cluster C (pink) and cluster G (green; solid lines intronic, dashed exonic. (c) Predicted percentage of canonical or non-canonical G-quadruplexes in DNMT1 eCLIP peaks from HeLa cells (Wang et al. 2025) versus pseudo-peaks in genes carrying no eCLIP peak.

